# Glucose restriction drives spatial re-organization of mevalonate metabolism and liquid-crystalline lipid droplet biogenesis

**DOI:** 10.1101/2020.08.29.273318

**Authors:** Sean Rogers, Hanaa Hariri, Long Gui, N. Ezgi Wood, Natalie Speer, Daniela Nicastro, W. Mike Henne

**Affiliations:** Department of Cell Biology, University of Texas Southwestern Medical Center, Dallas, Texas, 75024

## Abstract

Eukaryotes compartmentalize metabolic pathways into sub-cellular domains, but the role of inter-organelle contacts in organizing metabolic reactions remains poorly understood. Here, we show that in response to acute glucose restriction (AGR) yeast undergo metabolic remodeling of their mevalonate pathway that is spatially coordinated at nucleus-vacuole junctions (NVJs). The NVJ serves as a metabolic platform by selectively retaining HMG-CoA Reductases (HMGCRs), driving mevalonate pathway flux in an Upc2-dependent manner. AGR-induced HMGCR compartmentalization enhances mevalonate metabolism and sterol-ester biosynthesis that generates lipid droplets (LDs) with liquid-crystalline sub-architecture. Loss of NVJ-dependent HMGCR partitioning affects yeast growth, but can be bypassed by artificially multimerizing HMGCRs, indicating NVJ compartmentalization enhances mevalonate pathway flux by promoting HMGCR inter-enzyme associations. We propose a non-canonical mechanism regulating mevalonate metabolism via the spatial compartmentalization of rate-limiting HMGCR enzymes, and reveal that AGR creates LDs with remarkable phase transition properties.

**One Sentence Summary:** Spatial compartmentalization of HMG-CoA Reductases at ER-lysosome contacts modulates mevalonate pathway flux

## Introduction

The complexity of eukaryotic metabolism requires spatial organization, so membrane-bound and membraneless organelles compartmentalize enzymes into distinct sub-cellular regions. Enzyme partitioning into spatially defined domains is a cellular organizational principle observed in cytoplasmic assemblies (*1*), enzyme homo-polymers (*2*) and biomolecular condensates (*3, 4*). Such enzymatic assemblies, or metabolons, enhance or fine-tune metabolic flux, locally sequester toxic intermediates, or limit product shunting to competing pathways. Enzyme partitioning has also been exploited in synthetic bioengineering to artificially force enzymes into close proximity to drive local reactions or metabolic channeling through enzymatic cascades (*5*).

Inter-organelle contact sites also serve as platforms for enzymatic organization and lipid synthesis. In particular, the endoplasmic reticulum (ER) partitions lipogenic processes like non-vesicular lipid transport and organelle biogenesis at inter-organelle junctions (*6*–*8*). In yeast, ER-associated membranes (MAMs) harbor enhanced metabolic activity to support reactions such as the biosynthesis of phosphatidylethanolamine (*9*). More recent work has highlighted the role of ER-mitochondrial contacts defining distinct sub-domains within mitochondria that support the formation of multi-enzyme complexes that drive Coenzyme Q synthesis (*10, 11*). In yeast the nuclear envelope (continuous with the ER network) also makes extensive contact with the vacuole (equivalent to the lysosome), forming a nucleus-vacuole junction (NVJ) that acts as a metabolic platform to organize lipid transport, fatty acid synthesis, and lipid droplet (LD) biogenesis, particularly during metabolic stress cues (*7, 12, 13*). Despite these insights, few studies mechanistically dissect how metabolic cues regulate enzyme recruitment and compartmentalization at organelle-organelle contacts. Furthermore, how inter-organelle junctions enable enzyme spatial organization to fine-tune or enhance metabolic flux remains poorly described. Such inter-organelle crosstalk is particularly pertinent to understanding adaptive responses to stresses such as glucose starvation, which is characterized by drastic decreases in cellular ATP as well as changes in cytoplasmic pH and fluidity that alter macromolecular trafficking (*14, 15*).

Here, we utilize budding yeast as a genetically enabling model system to dissect the role of ER-lysosome contacts (e.g. the NVJ) as organizational platforms in mevalonate metabolism. We find that in response to acute glucose restriction (AGR), yeast actively partition and selectively retain HMG-CoA Reductase (HMGCR) enzymes at the NVJ in a Nvj1 and Upc2-dependent manner. This enzyme sub-compartmentalization enhances mevalonate pathway flux and promotes sterol-ester biosynthesis during AGR-induced metabolic remodeling. Failure to compartmentalize HMGCRs affects yeast growth, but can be rescued by the addition of exogenous mevalonate or the compartmentalization of HMGCRs via their artificial multimerization. Lastly, we find that AGR induces the metabolic remodeling of LDs, driving the formation of sterol-ester rich LDs that manifest striking phase transition properties due to lipolysis-dependent re-balancing of cellular neutral lipids.

## Results

### Yeast HMG-CoA Reductases (HMGCRs) inducibly partition at the NVJ in response to AGR

Given that ER-mediated contact sites in *Saccharomyces cerevisiae* act as lipogenic domains, we visually screened GFP-tagged neutral lipid metabolism enzymes for signs of compartmentalization at ER inter-organelle contact sites in yeast exposed to AGR. Note that AGR is operationally defined here as culturing yeast in synthetic complete (SC) media containing 2% glucose, then briefly centrifuging them and placing them into SC media containing 0.001% glucose. Screening revealed that endogenously GFP-tagged HMG-CoA Reductases (HMGCRs) Hmg1 and Hmg2, which localize throughout the ER network and particularly on the nuclear envelope (NE), enriched at regions where the nuclear surface was in contact with the vacuole (Fig 1A). This nucleus-vacuole interface was confirmed to be the NVJ as Hmg1-GFP co-localized with NVJ tether Nvj1-mRuby3 following AGR treatment for 4hrs (Fig 1B). As Hmg1 and Hmg2 are functionally redundant in mevalonate synthesis, a central metabolite that supplies diverse cellular pathways including sterol biogenesis, we chose to dissect the mechanisms underlying Hmg1 NVJ partitioning. To investigate whether the NVJ was required for Hmg1 partitioning, we imaged Hmg1-GFP in the absence of NVJ tethers Nvj1 and Vac8 (*16*), and found they were required for Hmg1-GFP partitioning during AGR (Fig 1C). Furthermore, time-lapse imaging revealed that Hmg1-GFP NVJ partitioning peaked after ∼4hrs of introducing yeast to AGR (Fig 1D-F). During this period, Hmg1-GFP was ∼15-times enriched at the NVJ relative to other NE regions. Collectively this suggests that AGR induces the time-dependent compartmentalization of Hmg1/2 at the NVJ in a Nvj1 and Vac8 dependent manner.

**Figure 1.**
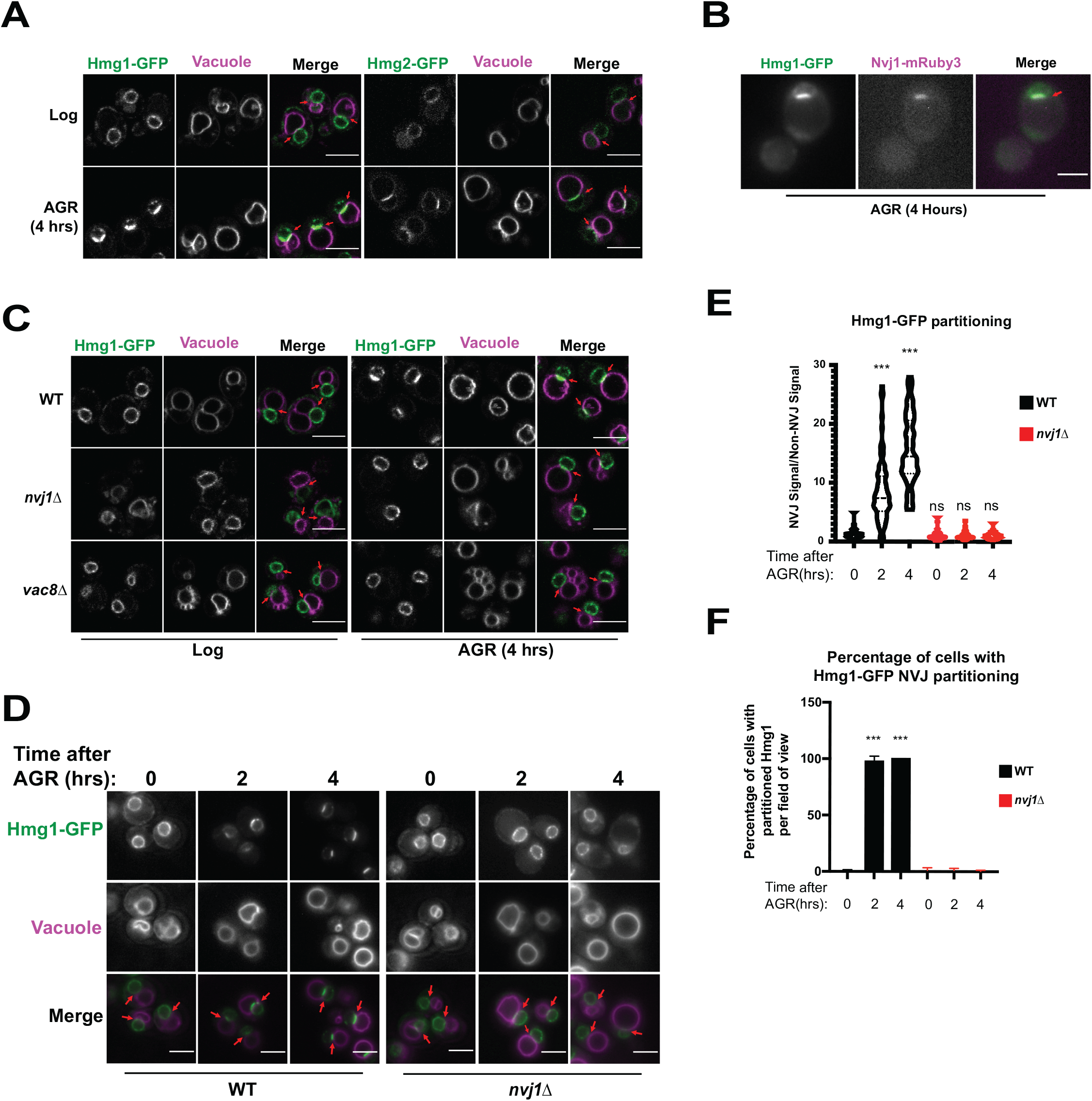
The HMG-CoA Reductases, Hmg1 and Hmg2, enrich at the NVJ in an Nvj1-dependent manner during AGR. (A) Confocal microscopy images of yeast expressing endogenously tagged Hmg1-GFP or Hmg2-GFP (green) grown in SC media with 2% glucose to an OD of 0.5 (Log), or in 0.001% glucose for 4hrs (AGR). Vacuoles (magenta) were visualized by staining with FM4-64. Red arrows represent the location of the Nucleus-Vacuole Junction (NVJ). Scale bars=5µm. (B) Epifluorescence microscopy images showing the overlap of Hmg1-GFP with Nvj1-mRuby3 after cells were exposed to four hours of AGR. Red arrows represent the location of the NVJ. Scale bars=5µm (C) Confocal microscopy images of yeast expressing endogenously tagged Hmg1-GFP (green) in wild type (WT), *nvj1*Δ, or *vac8*Δ yeast. Red arrows represent the location of NVJ. (D) Epifluorescence microscopy images of WT or *nvj1*Δ yeast expressing endogenously tagged Hmg1-GFP growing in 2% glucose to an OD of 0.5 (Log) or in 0.001% glucose for 2 or 4hrs (AGR). (E) Quantification of Hmg1 partitioning measured by line scan across the nuclear envelope (NE) and plotted as the ratio of Hmg1-GFP intensity at the NVJ relative to Hmg1-GFP intensity opposite the NVJ. Scale bars represent 5µm. (Brown-Forsyth and Welch ANOVA. N>51 cells. ***p-value<0.001). (F) Quantification of percentage of cells displaying Hmg1 NVJ partitioning per field of view. Any cell with partitioning >2.0 was considered as displaying Hmg1 NVJ partitioning. (Brown-Forsyth and Welch ANOVA. N>51 cells. ***p-value<0.001)

### NVJ compartmentalization of HMGCRs is independent of other mevalonate pathway enzymes and lipid trafficking proteins Osh1 and Ltc1

Inter-organelle contacts are implicated as domains coordinating the recruitment of supra-molecular enzyme complexes that constitute metabolic pathways. The HMGCRs are the rate-limiting enzymes of the mevalonate pathway that generates ergosterol, the cholesterol analog of yeast. To dissect whether other proteins along the mevalonate pathway also partition at the NVJ during AGR, we imaged 20 endogenously mNeonGreen (mNG)-tagged proteins involved in ergosterol biosynthesis under ambient and AGR stress, but surprisingly none detectably enriched at the NVJ (Fig 2A). As the NVJ has also been proposed to function in sterol transport between the vacuole and ER network, we also examined whether loss of the NVJ-resident lipid trafficking proteins Osh1 and Lam6/Ltc1 would impact Hmg1-GFP NVJ recruitment (*12, 17*). Hmg1-GFP efficiently partitioned at the NVJ following AGR in both *osh1*Δ and *lam6/ltc1*Δ yeast (Fig 2B). Collectively, this suggests that detectable Hmg1/2 NVJ enrichment is unique among enzymes involved in ergosterol metabolism, and is therefore unlikely to constitute a “classical” multi-enzyme metabolon.

**Figure 2.**
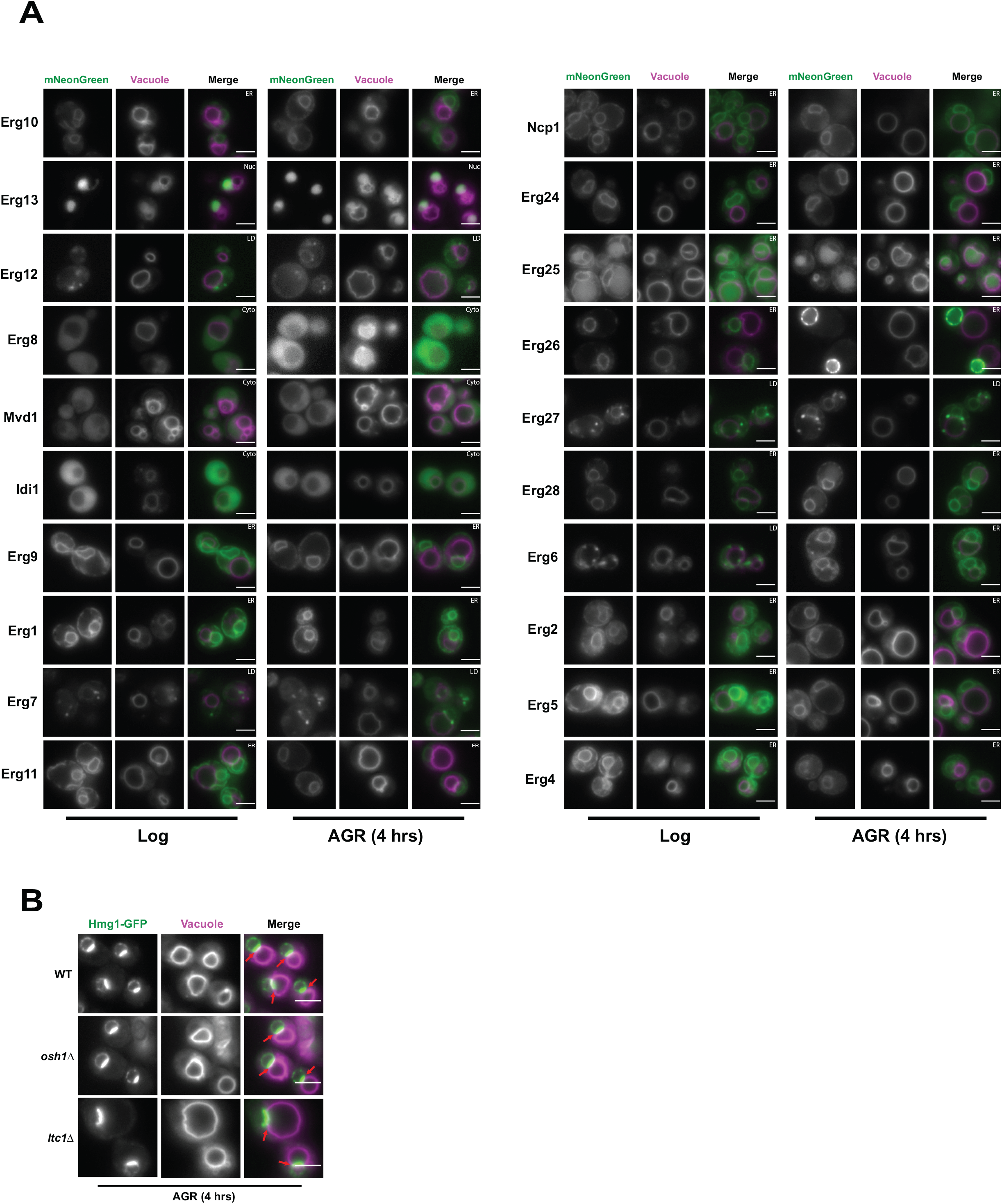
Hmg1 NVJ partitioning is specific among ergosterol biosynthesis proteins and independent of NVJ-associated sterol transport proteins. (A) Epifluorescence microscopy images of twenty ergosterol biosynthesis proteins endogenously tagged with mNeonGreen (mNg) imaged in log phase and after 4hrs of AGR. Cells were co-stained with FM4-64 (magenta) to visualize vacuoles. Cellular compartments occupied by each protein are designated in white text in the top right corner of each image. Scale bars=5µm. (B) Epifluorescence microscopy images of cells endogenously tagged with Hmg1-GFP in wild type (WT), *osh1*Δ, or *ltc1*Δ cells exposed to 4hrs of AGR. Scale bars=5µm.

### Hmg1-GFP is selectively retained at the NVJ in a Nvj1-dependent manner

Next, we investigated the mechanistic basis for NVJ compartmentalization of Hmg1-GFP. We began by interrogating whether the NVJ-partitioned and non-partitioned Hmg1-GFP pools were physically segregated along the NE. Using fluorescence recovery after photobleaching (FRAP), we determined that the non-partitioned Hmg1-GFP pool can enter the NVJ (Fig 3A,B); therefore, there is likely no diffusion barrier at the NVJ allowing selective entry of only specific pools of Hmg1-GFP protein. Next we interrogated whether Hmg1-GFP was selectively retained after entry into the NVJ region. Fluorescence loss in photobleaching (FLIP) revealed the average lifetime of NVJ-partitioned Hmg1-GFP was >100sec, compared to only ∼25sec for Hmg1-GFP along the rest of the NE (Fig 3C,D). Given that most of the Hmg1-GFP signal within the NVJ-partitioned population was not photobleached to completion, calculated halftimes may underestimate the actual halftime. Collectively, this indicates that upon AGR, Hmg1-GFP is recruited and selectively retained at the NVJ. Surprisingly, the halftime of Hmg1-GFP was also significantly increased during AGR in *nvj1*Δ cells, and further analysis is required to determine if this reduced mobility is specific to Hmg1-GFP or a general consequence of AGR.

**Figure 3.**
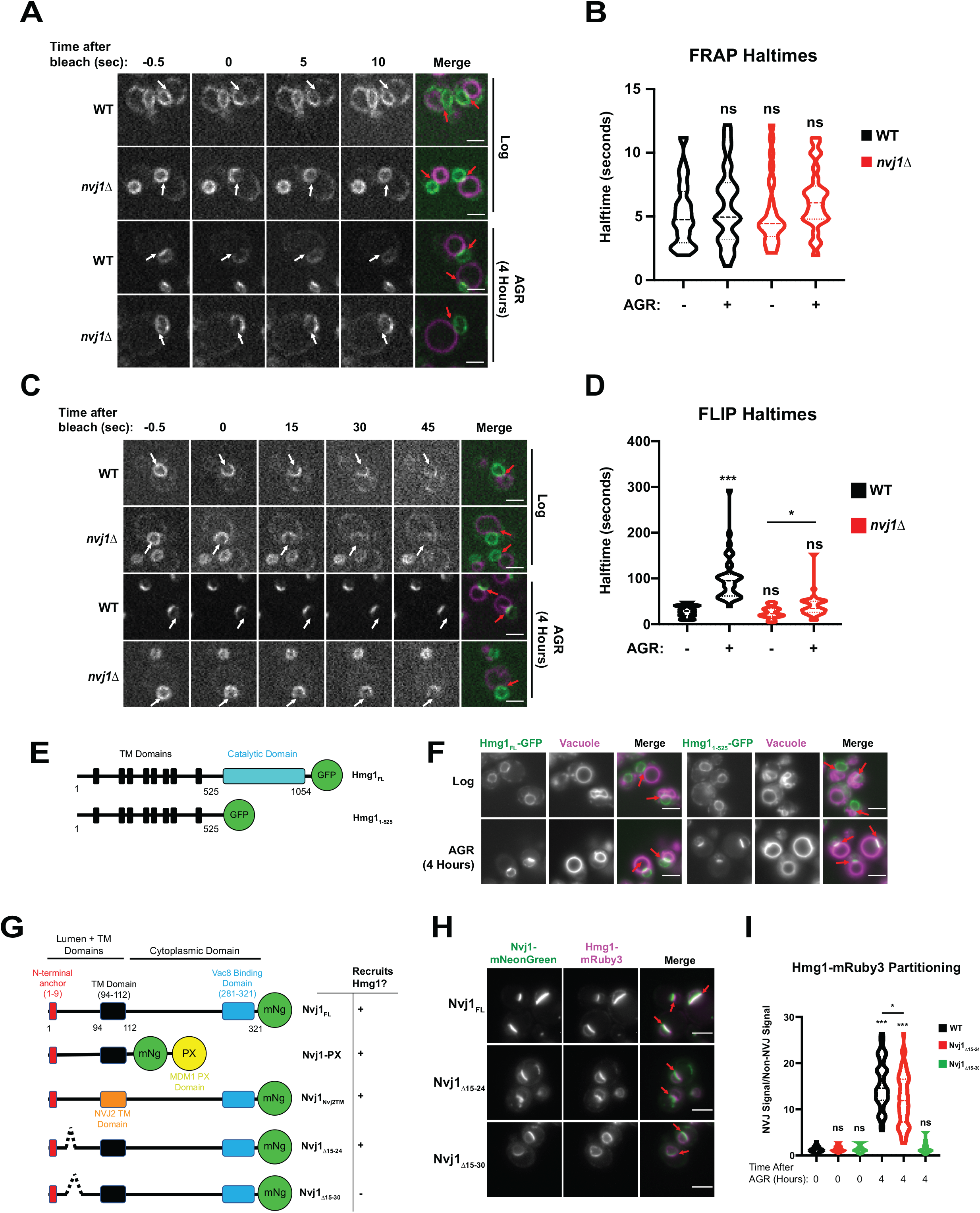
Hmg1 is selectively retained at the NVJ and requires a luminal domain of Nvj1. (A) Spinning disk confocal microscopy images from FRAP movies of yeast expressing endogenously tagged Hmg1-GFP and stained with FM4-64 (magenta). Portions of the NE corresponding to the NVJ were photobleached (white arrows) and recovery was monitored in both WT and *nvj1*Δ cells grown to log phase or after 4hrs of AGR. Red arrows represent the location of the NVJ. (B) Quantification of Hmg1-GFP FRAP halftimes after photobleaching. (Brown-Forsyth and Welch ANOVA. N>26 cells) (C) Spinning disk confocal microscopy images taken from FLIP movies. Conditions and strains were the same as in (A). One region of the NE that lies opposite the NVJ was photobleached every 5 seconds (white arrows) and Hmg1-GFP signal at the NVJ was monitored for loss of fluorescence. Red arrows represent the location of the NVJ (D) Quantification of FLIP halftimes. (Brown-Forsyth and Welch ANOVA. N>29 cells. *p-value>0.05 ***p-value<0.001). (E) Cartoon of Hmg1-GFP constructs endogenously expressed in yeast with both the cytosolic catalytic domain (blue) and TM/luminal domains (black) (Hmg1_FL_) or the TM/luminal domains alone (Hmg1_1-525_). (F) Epifluorescence microscopy images of yeast expressing endogenously tagged Hmg1_FL_-GFP or Hmg1_1-525_-GFP. (G) Cartoon of Nvj1-mNeonGreen (Ng) constructs expressed in *nvj1*Δ cells. Nvj1_FL_-mNg contains all domains of Nvj1 including the N-terminal luminal anchor (red), the TM domain (black), and the Vac8-binding domain (VBD) (blue). Chimeric constructs either replaced the VBD with a vacuole-binding PX domain (Nvj1_PX_) or replaced the endogenous TM domain with the TM domain of Nvj2 (Nvj1_NVJ2TM_). Images of these constructs can be found in Supplemental figure 1A. Truncations of Nvj1 removed either residues 15-24 (Nvj1_15-24Δ_) or 15-30 (Nvj1_15-30Δ_) (H) Epifluorescence microscopy images of yeast expressing truncation constructs depicted in (G). Cells were co-expressing endogenously tagged Hmg1-mRuby3 (magenta). Scale bars represent 5µm. (I) Quantification of Hmg1 partitioning at the NVJ for yeast expressing truncations of Nvj1-mNg and Hmg1-mRuby3. (Brown-Forsyth and Welch ANOVA. N>61 cells. *p-value<0.05 ***p-value<0.001)

Next, we dissected which regions of Hmg1 and Nvj1 were required for Hmg1-GFP NVJ retention. We generated truncated versions of GFP-tagged Hmg1 containing only its integral transmembrane (TM) region and lacking its cytoplasm-exposed catalytic domain. This Hmg1^1-525^-GFP truncation was sufficient to localize to the NVJ during AGR, indicating the integral membrane region of Hmg1 was sufficient for NVJ recruitment (Fig 3E,F). To dissect the Nvj1 regions required for Hmg1-GFP recruitment, we expressed mNG-tagged Nvj1 fragments in yeast co-expressing Hmg1-mRuby3 (Fig 3G). A Nvj1-mNG chimera lacking its cytoplasm-exposed region and containing a vacuole-binding PX domain in place of its Vac8-binding domain (Nvj1-PX) was sufficient to tether between the nucleus and vacuole, and was sufficient to partition Hmg1-mRuby3 at the NVJ during AGR, indicating the Nvj1 cytoplasmic domain is not required for Hmg1 recruitment (Supp Fig 1A). Similarly, a Nvj1-mNG chimera with its TM region replaced with the TM of Nvj2 (Nvj1_Nvj2TM_) also partitioned Hmg1-mRuby3 at the NVJ, indicating the TM and cytoplasmic regions are not required for Hmg1-mRuby3 recruitment (Supp Fig 1A). However, expression of a full length Nvj1-mNG lacking residues 15-30 (Nvj1_Δ15-30_) of its luminal region formed an NVJ contact but failed to recruit Hmg1-mRuby3. Re-addition of six residues to this construct (Nvj1_Δ15-24_) rescued Hmg1-mRuby3 recruitment, suggesting residues 25-30 of the Nvj1 luminal domain are required for Hmg1-mRuby3 NVJ partitioning (Fig 3G-I). Collectively, these data indicate that the ER embedded region of Hmg1 is recruited to the NVJ in a manner requiring the ER luminal Nvj1 region, and that Hmg1 partitioning is not dependent on the Hmg1 cytoplasmic catalytic domain.

### NVJ partitioning of Hmg1 is Upc2 dependent and correlates with sterol-ester biosynthesis

The NVJ was previously identified as a site for starvation-induced microautophagy of the nucleus (*18*), where NVJ-associated proteins accumulate and are subsequently turned over by the vacuole. HMGCRs are also known to undergo proteasomal degradation during specific metabolic cues. To determine if NVJ partitioning of Hmg1-GFP regulated Hmg1 protein abundance or turnover, we performed immunoblot analysis of Hmg1-GFP in log-phase and 4hrs AGR-treated cells. AGR increased steady-state Hmg1-GFP protein levels, indicating glucose restriction promoted Hmg1 protein accumulation (Fig 4A). Interestingly, *nvj1*Δ yeast displayed a more marked increase in Hmg1-GFP protein compared to wildtype. To determine whether Hmg1 protein accumulation was due to *de novo* protein synthesis, we co-treated yeast with the translational inhibitor cycloheximide during AGR. Indeed, cycloheximide suppressed AGR-induced Hmg1-GFP protein elevations, indicating that protein elevations were due primarily to new protein synthesis (Fig 4B). In line with this, treatment with the proteasome inhibitor MG132 during AGR had less effect on Hmg1 protein levels, suggesting that proteasomal turnover of Hmg1 was not a primary mechanism of protein abundance control during AGR (Fig 4B).

**Figure 4.**
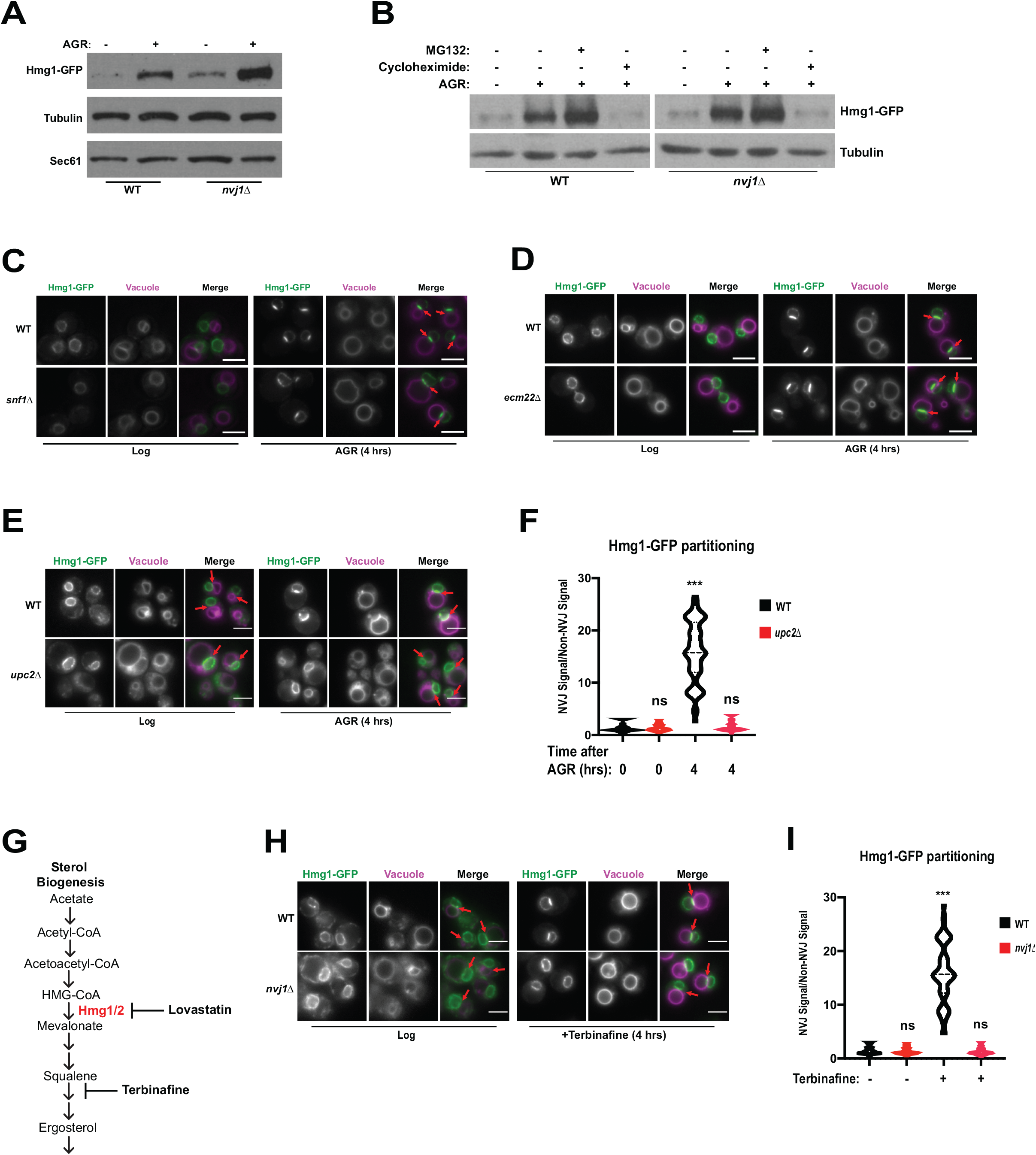
Hmg1 protein accumulates in *nvj1*Δ yeast via enhanced synthesis, which coincides with Upc2-dependent Hmg1 clustering at the NVJ. (A) Immunoblot of cells expressing endogenously-tagged Hmg1-GFP grown to log-phase or treated with AGR for 4hrs. Tubulin and Sec61 antibodies were used as loading controls. (B) Immunoblot of Hmg1-GFP expressing cells grown to log-phase or treated with AGR for four hours. AGR-treated cells were also co-treated with either 100µg/mL cycloheximide or 25µM MG132. Tubulin was used as a loading control. (C) Epifluorescence microscopy images of cells expressing endogenously tagged Hmg1-GFP in either a wild type (WT) or Snf1 knock-out (*snf1*Δ) background. (D) Epifluorescence microscopy images of cells expressing endogenously tagged Hmg1-GFP in either a wild type (WT) or *ecm22*Δ background. Scale bars=5µm. (E) Epifluorescence microscopy images of endogenously tagged Hmg1-GFP (green) with FM4-64 stained vacuoles (magenta) grown to log phase or after 4hrs of AGR in wild type (WT) and *upc2*Δ yeast. Scale bars=5µm. Red arrows indicate relative position of the NVJ. (F) Quantification of Hmg1-GFP partitioning at NVJ from images represented in (E). (Brown-Forsyth and Welch ANOVA. N>49 cells. ***p-value<0.001). (G) Cartoon representing abbreviated ergosterol biogenesis pathway. (H) Epifluorescence microscopy images of wild type and *nvj1*Δ cells expressing endogenously tagged Hmg1-GFP treated with 10µg/mL of terbinafine for four hours. Scale bars represent 5µm. (I) Quantification of Hmg1-GFP partitioning at the NVJ from images shown in (H). (Brown-Forsyth and Welch ANOVA. N>51 cells. ***p-value<0.001).

To further explore the metabolic cues governing AGR-induced Hmg1 synthesis and spatial partitioning, we monitored Hmg1-GFP localization in yeast lacking the major glucose-sensing kinase Snf1, the yeast AMPK homolog that regulates metabolic remodeling during changes in glucose availability (*19*). Surprisingly, *snf1*Δ yeast maintained AGR-induced Hmg1-GFP partitioning at the NVJ, indicating Hmg1-GFP recruitment to the NVJ was not dependent on Snf1/AMPK signaling (Fig 4C). We next examined whether Hmg1-GFP NVJ partitioning required the ergosterol-sensing transcription factor Upc2 that controls yeast sterol synthesis in a manner similar to mammalian SREBP signaling (*20*). Indeed *upc2*Δ yeast failed to partition Hmg1-GFP at the NVJ during AGR, and this was specific to *upc2*Δ, as Hmg1-GFP maintained NVJ partitioning in *ecm22*Δ yeast, an Upc2 paralog (Fig 4D-F). This implied that Hmg1-GFP NVJ partitioning correlated with alterations in cellular ergosterol levels. In line with this, Hmg1-GFP NVJ partitioning was also induced by treatment with the squalene epoxidase inhibitor terbinafine, which blocks *de novo* ergosterol biogenesis (Fig 4G-I).

We hypothesized that Hmg1-GFP NVJ enrichment correlated with elevations in cellular sterols. To test this, we conducted thin layer chromatography (TLC). Indeed, following 4hrs of AGR, yeast exhibited ∼30% more steady-state levels of sterol-esters (SE), while free ergosterol levels were unchanged (Fig 5A,B). Co-treatment with the HMGCR inhibitor lovastatin during AGR suppressed this SE elevation, indicating the elevated SE pool originated from *de novo* SE synthesis rather than esterification of pre-existing ergosterol. Altogether, this suggests that AGR induces an Upc2-dependent compartmentalization of *de novo* synthesized Hmg1 at the NVJ that correlates with elevated SE pools.

**Figure 5.**
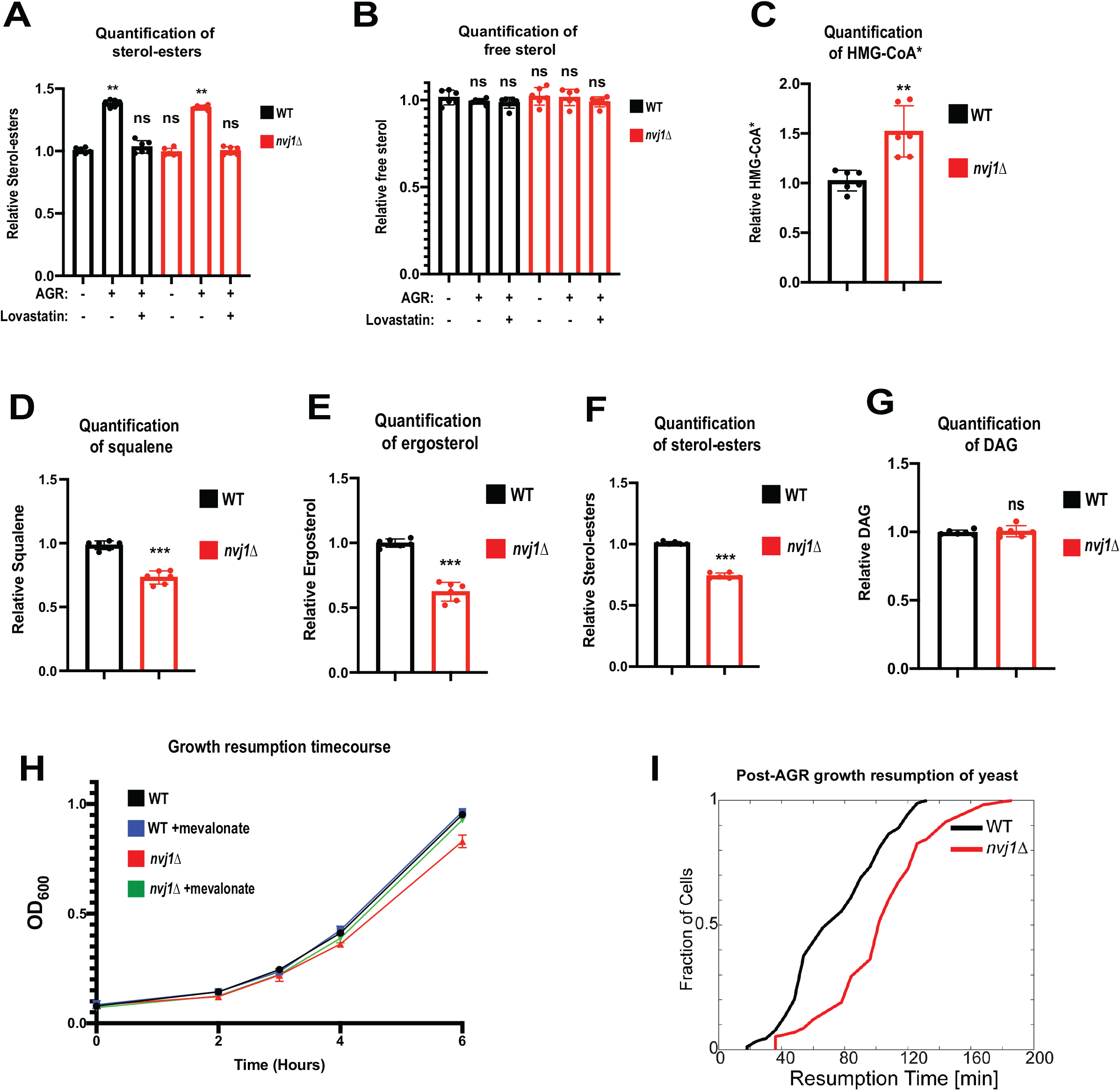
Hmg1 NVJ partitioning coincides with *de novo* sterol ester production and increased mevalonate pathway flux. (A) Quantification of SEs in wild type and *nvj1*Δ cells collected at log phase, after 4hrs of AGR, or after a 4hr co-treatment of AGR in the presence of 20µg/mL lovastatin. Neutral lipids were extracted, separated by TLC, and visualized using Cu(II) Sulfate spray and charring. Quantification performed by densitometry. (Brown-Forsyth and Welch ANOVA. N=6. **p-value<0.01). (B) Quantification of free sterols using the same strains and methodology as in (A). (Brown-Forsyth and Welch ANOVA. N=6. ***p-value<0.001). (C) Scintillation counting quantification of HMG-CoA* produced after a 15 minute pulse with ^14^C-acetate in intact WT or *nvj1*Δ cells grown for 4hrs in AGR. After quenching the radio-labeling reaction, soluble metabolites were isolated and endogenous HMG-CoA was converted into mevalonate using an *in vitro* HMGCR reaction. Mevalonate was subsequently separated and isolated by TLC, visualized by autoradiography, and quantified by scintillation counting. (Brown-Forsyth and Welch ANOVA. N=6. **p-value<0.01). (D-G) Quantifications of DAG and mevalonate-derived lipids. Radio-labeling was performed as described in (C). Lipids were extracted and separated by TLC and visualized by autoradiography. Squalene and SE bands were quantified by scintillation counting. Ergosterol and DAG bands were quantified by densitometry. (Brown-Forsyth and Welch ANOVA. N=6. ***p-value<0.001). (H) Growth curves of WT and *nvj1*Δ cells. Cells were treated with AGR for 10 hrs in the absence or presence of 10µg/mL mevalonate, and subsequently diluted to OD_600_=0.1 in SC media containing 2% glucose lacking mevalonate. OD_600_ measurements were taken every hour following dilution in fresh glucose-containing media. (I) Quantification of growth resumption in WT and *nvj1*Δ cells after ten hours in AGR. Resumption of growth was scored using single-cell time-lapse imaging (see Methods). Cells lacking Nvj1 exhibit approximately a 30-minute delay in growth resumption after re-introduction to glucose.

### Loss of HMGCR NVJ partitioning affects mevalonate flux and yeast growth

Given that *nvj1*Δ yeast accumulate more Hmg1-GFP protein than wildtype cells (Fig 4A), we were surprised to find that *nvj1*Δ yeast produce similar steady-state levels of SEs following AGR (Fig 5A). This implied that HMGCR enzymes in *nvj1*Δ yeast could be catalytically less efficient. If true, we would expect: 1) accumulation of the HMGCR substrate HMG-CoA, and 2) a decrease in downstream mevalonate pathway products such as squalene, ergosterol, and SEs. To interrogate whether Hmg1 spatial compartmentalization influenced Hmg1 enzymatic activity and/or mevalonate pathway flux, we used ^14^C-acetate pulse-radiolabeling to monitor these mevalonate pathway components. Indeed, *nvj1*Δ yeast exhibited significantly elevated ^14^C-labeled HMG-CoA after a 15-minute radio-pulse, and contained significantly less ^14^C-labeled squalene, ergosterol, and SE (Fig 5C-F, Supp Fig 2A). However, ^14^C-diacylglycerol (DAG) was not significantly altered, suggesting these alterations were specific to mevalonate synthesis and pathway flux (Fig 5G).

To investigate whether these alterations in mevalonate metabolism affected yeast fitness or growth, we monitored yeast growth in glucose-containing SC media following a 10hr AGR treatment. Indeed, *nvj1*Δ yeast displayed slower growth in batch cultures compared to wildtype, and this was rescued by addition of exogenous mevalonate, suggesting the growth defect was attributed to defects in the mevalonate pathway (Fig 5H). To dissect the nature of this delayed cell growth, we conducted single-cell time-lapse imaging during 10hrs of AGR, and a subsequent phase when yeast were re-supplied with SC-media containing 2% glucose. This revealed that *nvj1*Δ yeast displayed a significant delay in growth resumption following AGR stress when re-exposed to glucose-containing media (Fig 5I). In line with this, doubling times of cells were not affected by either loss of Nvj1 or addition of mevalonate following growth resumption (Supp Fig 2B). Together, these findings indicate that *nvj1*Δ yeast unable to spatially compartmentalize Hmg1 manifest alterations in mevalonate pathway flux, and that *nvj1*Δ yeast manifest growth delay defects following AGR that can be rescued by exogenous mevalonate addition.

### Uncoupling Hmg1 spatial compartmentalization from protein abundance reveals a role for NVJ partitioning in mevalonate flux

The AGR-induced changes in Hmg1 protein abundance made it difficult to specifically dissect the role of Hmg1 spatial compartmentalization in mevalonate metabolism. To mechanistically dissect this compartmentalization and uncouple it from protein level, we generated yeast strains lacking endogenous Hmg1 and Hmg2 and expressing Hmg1-GFP from a non-native *ADH* promoter (Fig 6A). This new strain (*hmg1*Δ*hmg2*Δ ADHpr:Hmg1-GFP) maintained similar Hmg1-GFP protein levels with and without AGR treatment, and in the presence or absence of Nvj1 (Fig 6B). Critically, these yeast still partitioned Hmg1-GFP at the NVJ during AGR in an Nvj1-dependent manner, indicating we had uncoupled Hmg1 protein levels from Hmg1-GFP spatial compartmentalization (Fig 6C).

**Figure 6.**
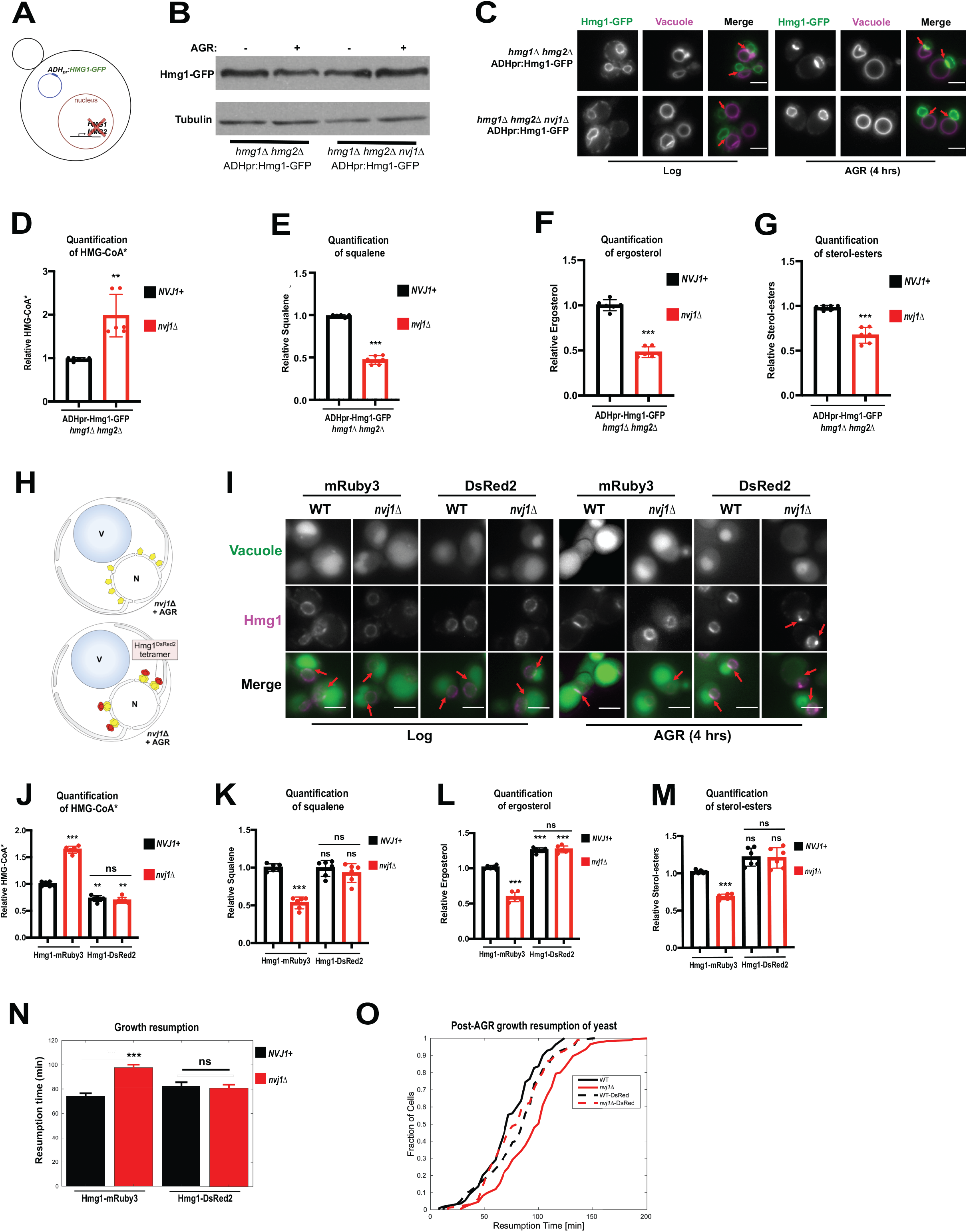
AGR-induced compartmentalization of Hmg1 increases mevalonate pathway flux through inter-enzyme associations. (A) Cartoon representation of *hmg1*Δ*hmg2*Δ ADHpr:Hmg1-GFP strains. Strains were generated by removal of endogenous Hmg1 and Hmg2, and replacing with ADH promoter-driven Hmg1-GFP (ADHpr:Hmg1-GFP). (B) Immunoblot of Hmg1-GFP isolated from ADHpr:Hmg1-GFP strains before or after treatment with AGR. Tubulin was used as a loading control. (C) Epifluorescence imaging of ADHpr:Hmg1-GFP strains showing that Hmg1 NVJ partitioning can be uncoupled from endogenous protein expression. Red arrows indicate relative position of the NVJ. Scale bars = 0.5µm. (D) Quantification of HMG-CoA* from yeast expressing ADHpr:Hmg1-GFP in wildtype (*NVJ1*+) or *nvj1*Δ yeast. Growth and radiolabeling was performed as described above. (Brown-Forsyth and Welch ANOVA. N=6. **p-value<0.01). (E-G) Quantification of squalene, ergosterol, and SE produced from a 15 minutes radio-labeling pulse in ADHpr:Hmg1-GFP strains. (Brown-Forsyth and Welch ANOVA. N=6. ***p-value<0.001). (H) Cartoon representation of Hmg1 artificial tetramerization by DsRed2 tagging in *nvj1*Δ cells. (I) Epifluorescence imaging of yeast expressing endogenously tagged Hmg1-mRuby3 or Hmg1-DsRed2 (magenta) in log-phase and after four hours of AGR treatment. Vacuoles (green) were stained by incubating yeast with 5µg/mL CMAC dye for two hours prior to imaging. Red arrows in merged imagew indicate the NVJ. Red arrows in the Hmg1-DsRed2 *nvj1*Δ AGR-treated greyscale image indicates prominent Hmg1-positive foci on the NE. Scale bars=0.5µm. (J) Scintillation counting quantification of HMG-CoA* from yeast expressing endogenously tagged monomeric Hmg1-mRuby3 or tetrameric Hmg1-DsRed2 in wildtype (*NVJ1*+) or *nvj1*Δ yeast. Quantification performed as in (D). (Brown-Forsyth and Welch ANOVA. N=6. **p-value<0.01 ***p-value<0.001). (K-M) Quantification of squalene, ergosterol, and SE produced in Hmg1-mRuby3 or Hmg1-DsRed2 yeast. Radio-labeling and quantifications performed as in (E-G). (Brown-Forsyth and Welch ANOVA. N=6. ***p-value<0.001). (N) Average growth resumption times measured by single-cell time-lapse microscopy. (WT: N=110, nvj1Δ: N=175, WT Hmg1-DsRed2: N=94, nvj1Δ Hmg1-DsRed2: N=96). (**p<0.001, Kolmogorov-Smirnov test). (O) Empirical cumulative distribution function of the resumption times. (WT: N=110, nvj1Δ: N=175, WT-DsRed: N=94, nvj1Δ-DsRed: N=96).

Next, we dissected whether loss of NVJ spatial compartmentalization in this ADHpr:Hmg1-GFP yeast impacted mevalonate pathway flux. Remarkably, radio-pulse analysis revealed that *nvj1*Δ yeast with ADHpr:Hmg1-GFP displayed exacerbated alterations in mevalonate pathway flux, exhibiting elevated ^14^C-labeled HMG-CoA levels and decreased ^14^C-squalene, ergosterol, and SEs (Fig 6D-G, Supp Fig 3A). Once again, ^14^C-DAG levels were unaffected, indicating that *nvj1*Δ yeast manifested specific defects in mevalonate metabolism and not other ER-associated lipid metabolic pathways (Supp Fig 3B). This suggests that Hmg1-GFP NVJ spatial partitioning independent of Hmg1 protein levels promotes mevalonate pathway flux.

### Loss of Hmg1 NVJ partitioning can be bypassed by compartmentalizing Hmg1 via artificial multimerization

Bioengineering studies indicate that HMGCR catalytic domains exhibit enhanced catalytic activity when forced into close proximities via multi-valent flexible scaffolds (*5*). Similarly, human HMGCR requires tetramerization for catalytic activity (*21*). We hypothesized that Hmg1 NVJ partitioning may enhance enzymatic activity by promoting close physical associations between HMGCR enzymes. To test this, we fused Hmg1 to a constitutively tetrameric fluorescent protein, DsRed2, and determined whether this artificial multimerization could bypass the loss of NVJ-mediated compartmentalization by comparing it to Hmg1 tagged with a monomeric fluorescent protein, mRuby3 (Fig 6H). Indeed, Hmg1-DsRed2 foci appeared on the NE during AGR, consistent with stabilized Hmg1 multimers (Fig 6I). As expected, *nvj1*Δ cells tagged with mRuby3 still manifested elevated HMG-CoA levels and reduced squalene, ergosterol, and SE pools associated with NVJ loss. Strikingly, these perturbations were rescued in the *nvj1*Δ Hmg1-DsRed2 strain, closely mirroring wildtype levels of Hmg-CoA, squalene, ergosterol, and SE (Fig 6J-M, Supp Fig 4A,B). This indicated that artificial multimerization of Hmg1 could bypass the loss of NVJ-mediated Hmg1 compartmentalization.

Next, we interrogated whether Hmg1-DsRed2 tagging would rescue the growth resumption delay observed for *nvj1*Δ yeast following AGR stress. As expected, growth resumption delay was observed in *nvj1*Δ yeast expressing Hmg1-mRuby3. Remarkably, expression of Hmg1-DsRed2 in *nvj1*Δ cells rescued this defect (Fig 6N, 6O). Collectively, this is consistent with a model where NVJ-dependent Hmg1 spatial compartmentalization via NVJ partitioning or artificial multimerization enhances mevalonate pathway flux.

### Acute glucose restriction promotes the formation of SE phase transitions within LDs

Given that AGR significantly increased cellular SE levels, and the NVJ is a site for the biogenesis of lipid droplets (LDs) that store SEs (*7*), we visualized LDs during AGR using high resolution cryo-focused ion beam-scanning electron microscopy (cryoFIB-SEM). Imaging revealed that non-AGR treated yeast contained LDs with homogenous lipid interiors. However, ∼77% of LDs observed in 4hrs AGR manifested LDs with multi-layered concentric rings (Fig 7A-F). The concentric ring layers were separated by ∼3.3nm, appearing similar to liquid-crystalline (LC) SEs observed in HeLa cells following mitotic arrest (*22*) (Fig 7E,G). Previous *in vitro* work of similar LC-layered LDs (LCL-LDs) demonstrated that specific SE-to-TG ratios were required to maintain the SE phase transitions within LCL-LDs (*23*). In line with this, TLC revealed that AGR-induced a ∼50% decrease in cellular TG, indicating TG levels were reduced during periods of SE elevation during AGR (Fig 7H).

**Figure 7.**
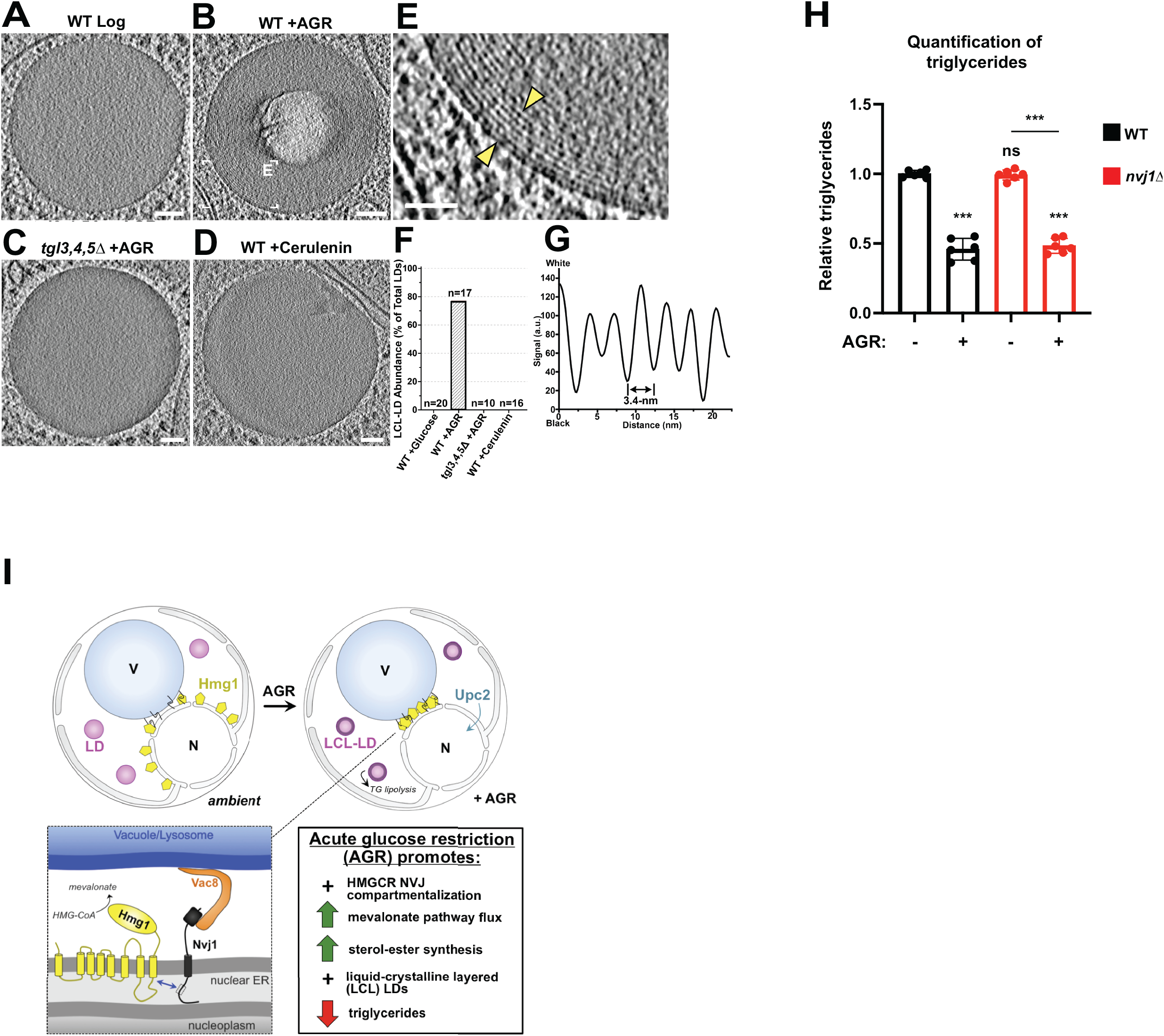
AGR promotes TG lipolysis and formation of LCL-LDs. (A-G) 1.1 nm thick tomographic slices from cryo-FIB-milled and cryo-ET reconstructed WT yeast cells grown with glucose (A), in AGR (B), *tgl3,4,5*Δ mutant yeast cell in AGR (C), and WT yeast grown in glucose treated with 10µg/mL cerulenin to induce TG lipolysis (D). Liquid-crystalline layered (LCL) LDs were only observed in WT+AGR. The boxed area in B is magnified in (E). (F) Abundance of LCL-LDs in WT yeast cells grown with glucose, WT in AGR, *tgl3,4*,5Δ in AGR, and WT treated with cerulenin. (G) Line plot displaying the gray values across the region of the LC layers (between yellow arrowheads in E) shows a 3.4-nm interval between adjacent layers. Scale bars=50 nm (A-D) and 25 nm (E). (H) Quantification of TG in log-phase and AGR-treated cells. (I) Cartoon representation of Nvj1 and Upc2-mediated Hmg1 partitioning at the NVJ. Spatial partitioning of Hmg1 at the NVJ coincides with increased production of mevalonate-derived metabolites.

We investigated whether TG lipolysis was required for LCL-LD formation. Indeed, yeast lacking the three TG lipases (*tgl3,4,5*Δ) did not exhibit LCL-LDs following 4hrs AGR (Fig 7C,F). However, stimulating lipolysis by culturing yeast with cerulenin was not sufficient to create LCL-LDs, implying lipolysis alone is not sufficient for LCL-LD formation (Fig 7D,F). Finally, we noted that LCL-LDs formed during AGR became significantly more prone to radiation damage during SEM imaging, which was not observed for homogenous LDs (Supp Fig 5A-J). This suggested that the cores of LCL-LDs may undergo substantial biochemical re-organization during AGR. Collectively, these results indicate that AGR stress promotes LCL-LD formation that involves *de novo* SE production as well as Tgl-dependent TG lipolysis.

## Discussion

Acute changes in nutrient availability trigger metabolic remodeling that is characterized by alterations in metabolic flux and metabolite supply and demand. How cells spatially and temporally coordinate this remodeling remains poorly characterized, yet critical to our understanding to cell adaptation and survival. Changes in nutrient availability can drive the sub-cellular re-distribution of enzymes within cells, as well as the formation of enzyme assemblies or complexes that promote or fine-tune metabolic pathways. Here, we present evidence that AGR in yeast enhances mevalonate pathway flux, and propose the mechanism underlying this requires the spatial compartmentalization of the rate-limiting HMGCR enzymes at the yeast NVJ (Fig 7I). Through time-lapse and FRAP/FLIP-based imaging, we find that Hmg1 is recruited and selectively retained at the NVJ in an Nvj1-dependent manner. This retention correlates with elevated *de novo* SE synthesis, as well as the Upc2 sterol-sensing transcription factor. Remarkably *nvj1*Δ yeast manifest alterations in mevalonate pathway flux and growth defects when faced with glucose restriction, which can be rescued by the addition of exogenous mevalonate (the HMGCR enzymatic product) or via the compartmentalization of Hmg1 via its artificial multimerization. To more fully dissect the role of HMGCR spatial compartmentalization on mevalonate metabolism, we generated an artificial yeast strain that maintains constant Hmg1 protein levels and inducibly accumulates at the NVJ, thus functionally uncoupling Hmg1 protein abundance from NVJ spatial compartmentalization. Remarkably, this strain exhibits defects in mevalonate pathway flux when Hmg1 cannot be NVJ partitioned, underscoring the role of Hmg1 spatial compartmentalization in fine-tuning mevalonate metabolism. Collectively, we present a model where AGR-induced metabolic remodeling of mevalonate metabolism is spatially coordinated at the NVJ via selective retention of HMGCRs.

In addition to inducing the compartmentalization of HMGCRs at the NVJ, yeast glucose starvation has also been shown to induce the formation of cytoplasmic enzyme assemblies involved in nucleotide (Ura7), amino acid (Gly1), and lipid (Acs1) metabolism (*15*). These cytoplasmic assemblies are thought to reduce enzymatic activity and promote the transition into cellular dormancy. In contrast, other protein assemblies promote enzymatic activity. In the liver, the protein Mig12 binds to cytosolic acetyl-CoA carboxylases (Acc) and promotes their polymerization and catalytic activity, which elevates fatty acid synthesis and eventual triglyceride accumulation in hepatocytes (*24*). Our work provides evidence that HMGCRs form similar multi-enzyme assemblies at yeast ER-lysosome contact sites, implicating a role for inter-organelle junctions in the spatial coordination of metabolically-induced enzymatic assemblies.

Our work also reveals that AGR induces the remodeling of cellular neutral lipid pools and promotes lipid phase transitions within LDs. This neutral lipid remodeling is coordinated via an elevation of SEs from *de novo* synthesis and con-commitment TG decrease via lipolysis. As LDs are the primary neutral lipid storage organelles of eukaryotes, we utilized cryoFIB-SEM to visualize LDs under ambient and AGR growth conditions. Strikingly, AGR-induced phase transitions can be directly visualized, giving rise to onion-like layering at the periphery of the LD interior. Similar phase transitions have been observed in mammalian cells faced with mitotic arrest or starvation, indicating LC transitions within LDs are highly conserved (*22*). We find that LCL-LD formation requires TG lipases, indicating that SE phase transitions are likely promoted by lowering LD-resident TG levels via Tgl-dependent lipolysis. In line with this, SE LC transition is inhibited by TG *in vitro* (*15, 17*). The LC layering of the LD surface likely induces changes in the LD surface proteome, and how these alterations affect LD function and inter-organelle interactions will be the focus of future studies. The LC interiors of LCL-LDs are conceptually similar to SE-rich LDL particles and atherosclerotic plaques, two biomedically relevant macromolecules characterized by LC SE phase transitions (*25, 26*). Further insights into how lipolysis and SE synthesis contribute to LCL-LD biogenesis and SE phase transitions may provide unexpected insights into cellular sterol metabolism and human cardiovascular diseases like atherosclerosis.

## Materials and Methods

### Strains, plasmids, and yeast growth conditions

W303 (leu2-3,112 trp1-1 can1-100 ura3-1 ade2-1 his3-11,15) was used as the wild type parental strain for all experiments and cloning in this study. All strains and plasmids can be found in tables S1 and S2, respectively. Deletion of endogenous genes, C-terminal tagging at endogenous loci, and transformation of plasmids was accomplished by the traditional lithium acetate method. Endogenous knockins and knockouts were validated by genomic PCR, and plasmids were validated by sequencing. Plasmids generated for this study were created using RepliQa HiFi assembly (Quantabio cat. 95190) following the manufacturer’s protocol. For cloning into pRS305 plasmids, destination vectors were cut with XhoI and SacI enzymes, while linearization of pFa6a plasmids was accomplished by digestion with AscI and PacI enzymes.

Synthetic complete media supplemented with amino acids was used for all experiments, except where leucine was omitted from the media to accommodate growth of strains carrying pRS305 plasmids. All yeast were grown in 30C incubators shaking at 210RPM. Log phase yeast were grown to an OD_600_ of 0.5 in the presence of 2% dextrose. AGR-treated yeast were grown to OD_600_ of 0.5 in the presence of 2% dextrose, collected, washed with dextrose-free media, and resuspended in media containing 0.001% dextrose for the indicated time period. For experiments where TG lipolysis was induced, cells were grown to stationary phase over 24 hours and then diluted to an OD_600_ of 0.5 in the presence of 2% dextrose and 10µg/mL Cerulenin (Sigma C2389). Cells were incubated with cerulenin for four hours prior to collection. Where indicated, Terbinafine (Sigma T8826), Lovastatin (Sigma 1370600), Cycloheximide (Sigma C7698), or MG132 (Sigma M7449) were added to the media at the beginning of AGR treatment to final concentrations of 10µg/mL, 20µg/mL, 100µg/mL, and 25µM, respectively.

### Fluorescence microscopy

For confocal microscopy, cells were grown as described above, collected by centrifugation at 3,000xg for two minutes, and resuspended in glucose-free media at approximately one one-hundredth of original volume. All images were taken as single slices at approximately mid-plane using a Zeiss LSM880 inverted laser scanning confocal microscope equipped with Zen software. Images were taken with a 63x oil objective NA=1.4 at room temperature. Prior to imaging, cells were incubated for three hours with 0.5µg/mL FM4-64 dye (Invitrogen T13320) to visualize vacuoles.

For epifluorescence microscopy, cells were grown, stained, and collected as described above. Vacuoles were also stained with 5µg/mL CMAC (ThermoFisher C2110) dye for two hours prior to imaging, where indicated. Imaging was performed on an EVOS FL Cell Imaging System at room temperature. Hmg1 NVJ partitioning was quantified using Fiji software. For quantification, RGB images were converted to 16-bit and a background subtraction was performed by subtracting original images by a duplicate ‘Gaussian blur’ filtered image (sigma (radius)=5.0). Five-pixel line scans were taken across the nuclear envelope toward the NVJ, and the ‘plot profile’ function was used in Fiji to produce a fluorescence histogram of nuclear envelope signal. The sum area under each curve was calculated and plotted as the ratio of fluorescence intensity of NVJ-associated signal by fluorescence intensity of Non-NVJ associated NE signal.

### FRAP and FLIP analysis

Yeast used for FRAP and FLIP were grown and collected as described above, and imaging was conducted for one hour after collection. Photobleaching movies were taken on an Andor spinning disk confocal microscope through a 63x oil objective (NA=1.4). The microscope is equipped with an Andor Ultra EMCCD and Metamorph software. For FRAP measurements, a single circular ROI of 0.77µm^2^ that corresponds to the NVJ was selected and bleached with a 408nm laser at 100% power and 100ms dwell time. One image was taken before the bleach, and subsequent images were taken every 500ms for a total movie length of 25 seconds. For FLIP measurements, single circular ROIs of 0.77µm^2^ were selected, taking care to select an area of the NE that was furthest from the NVJ. Each bleaching cycle consisted of a pre-bleach image, a single bleach with a 408nm laser with a 100ms dwell time, and four post-bleach images taken 500ms apart. Each movie captured a total of 50 bleach cycles, which corresponds to 300 second movies. Fiji software was used to quantify bleaching curves and halftimes. Pre-processing of images included background subtraction as described above and 3D Gaussian smoothing (sigma=0.5). FRAP quantification was performed using the double normalization method as previously described (*27*). Briefly, intensity was measured for all time points in an ROI corresponding to the bleached region (I_frap_) and an ROI corresponding to the whole cell (I_whole-cell_), and normalized intensities were generated for each timepoint (I_normalized_(t)) using equation S1:

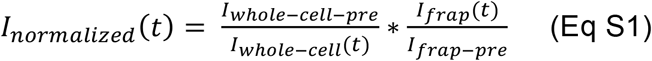

In the equation above, the ‘pre’ subtext indicates the timepoint preceding ROI bleaching. Intensity recovery curves were created for each movie by further normalizing values such that pre-bleach intensities were set to 1 and post-bleach intensity of an ROI was set to 0. Full normalization was accomplished using equation S2 followed by subtracting the normalized I_frap-bleach_ value from all timepoints.

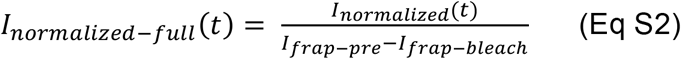

In equation S2, I_frap-bleach_ indicates the intensity of the photobleached ROI at the time of photobleaching. Halftimes were calculated from individual fluorescence recovery curves using Graphpad Prism 8 software and fitting the data to a one-phase exponential association. Pre-processing for FLIP images was the same as for FRAP images. FLIP movies were quantified using Fiji to monitor the intensity of NVJ-associated signal over time. Intensity measurements were normalized using equation S3.

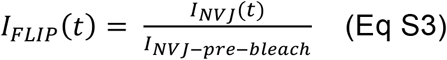

In equation S3, I_FLIP_(t) is the relative fluorescence at the NVJ at time t, I_NVJ_(t) is the raw intensity value at the NVJ at time t, and I_NVJ-pre-bleach_ is the intensity at the NVJ before bleaching occurred. Halftimes were calculated from FLIP decay curves, which were generated in Graphpad Prism 8 software by fitting the data to a one-phase exponential decay.

### Lipid extraction and thin layer chromatography

For lipid extraction, approximately 50OD units of cells were collected for each sample, and pellet wet weight was normalized and noted prior to extraction. Lipid extraction was performed using a modified Folch method (*28*). Briefly, cell pellets were resuspended in MilliQ water with glass beads and lysed by three one-minute cycles on a bead beater. Chloroform and methanol were added to the lysate to achieve a 2:1:1 chloroform:methanol:water ratio. Samples were vortexed, centrifuged to separate the organic and aqueous phases, and the organic phase was collected. Extraction was repeated a total of three times. Prior to thin layer chromatography, lipid samples were dried under a stream of argon gas and resuspended in 1:1 chloroform:methanol to a final concentration corresponding to 4µL of solvent per 1mg cell pellet wet weight. Isolated lipids were spotted onto heated glass-backed silica gel 60 plates (Millipore Sigma 1057210001), and neutral lipids were separated in a mobile phase of 80:20:1 hexane:diethyl ether:glacial acetic acid. TLC bands were visualized by spraying dried plates with cupric acetate in 8% phosphoric acid and baking at 140C for an hour. To quantify TLC bands, all plates were run with an internal neutral lipid standard. Densitometry of bands was performed in Fiji.

### Batch culture growth curves

Cells were treated with AGR for ten hours, as described above. Some samples were also co-treated with 10µg/mL mevalonate (Sigma 50838) at the beginning of AGR treatment. After a ten hour treatment with AGR, cultures were diluted to an OD_600_ of 0.1 in SC media containing 2% glucose. The OD_600_ of cultures was measured each hour and plotted in Prism 8 software.

### Single cell time-lapse microscopy

Cells were imaged using a Zeiss Observer Z1 microscope equipped with automated hardware focus, motorized stage, temperature control, a Zeiss EC Plan-Apochromat 63X 1.4 or 40X 1.3 oil immersion objectives, and an AxioCam HRm Rev 3 camera. Exposure times for experiments shown in Supp Fig 5B: Phase contrast 20ms, Whi5-mKoκ 150 ms, Erg6-mTFP1 20 ms, Vma1-mNeptune2.5 75ms, Nvj1-mRuby3 75ms, Msn2-mNeonGreen 40ms. Exposure times for experiments shown in Fig 4 and Supp Fig 7: Phase contrast: 40 ms, Hmg1-mRuby3 or Hmg1-DsRed2: 100 ms.

All experiments were performed with a Y04C Cellasic microfluidic device (http://www.cellasic.com/) using 1 psi flow rate. Images are taken every 6 minutes. Prior to loading into the microfluidics chamber, cells were sonicated and mixed with 50µL SCD media to achieve an OD_600_ of approximately 0.1. In the chamber, cells are first grown for 2hrs in SCD. Next, they are exposed to acute glucose restriction (AGR) by switching to SC for 10 hrs which is followed by glucose replenishment by 4 hrs SCD. To determine time of growth resumption, cells are segmented and tracked as described previously (*29, 30*). Next, the time of resumption is annotated semi-automatically using a custom MATLAB software by determining the time of bud growth or new bud emergence during the four-hour glucose replenishment following AGR.

### Immunoblotting

Approximately 50OD units of cells was collected for protein extraction. Prior to protein extraction, cell pellet wet weights were normalized. Protein extraction was accomplished by precipitating proteins with 20% TCA for thirty minutes on ice, followed by three washes of the pellet with cold 100% acetone. The protein pellet was dried for fifteen minutes in a speed-vac to remove residual acetone, and all pellets were resuspended in 2x SDS sample buffer (65.8mM Tris-HCl, pH 6.8; 2% SDS; 25% glycerol; 0.01% bromophenol blue). Resuspended protein samples were heated at 70C for ten minutes prior to being loaded onto a homemade 4-15% polyacrylamide gel and separated by electrophoresis. Proteins were transferred to a 0.45µm nitrocellulose membrane in Towbin SDS transfer buffer using a Criterion tank blotter with plate electrodes (BioRad 1704070). Membranes were blocked with 5% milk and primary antibodies were allowed to bind overnight at 4C. Hmg1-GFP was detected using a rabbit polyclonal antibody to GFP (Abcam ab290; 1:10,000 dilution). Rat monoclonal antibody to tubulin (Abcam ab6160; 1:15,000 dilution) and rabbit polyclonal antibody to Sec61 (Jonathan Friedman lab; 1:5,000 dilution) were used as loading controls. Immunoblots were developed by binding HRP-conjugated anti-rabbit IgG (Sigma A0545; 1:10,000) or anti-rat IgG (Abcam ab97057; 1:10,000) antibodies to the membrane for one hour followed by developing in ECL (BioRad 1705061). Signal was captured by x-ray film.

### Cryo-sample preparation and Cryo-FIB milling

4 µl of the cultured yeast cells (OD600 of 0.4) were added to a glow-discharged (30 seconds at -30 mA) copper R2/2 holey carbon grid (Quantifoil Micro Tools GmbH, Jena, Germany), then the grid was rapidly plunge frozen in liquid ethane using a homemade plunge freezer and stored in liquid nitrogen until used. Cryo-FIB milling was performed as previously described (*31*). Briefly, grids were mounted in notched cryo-FIB Autogrids (Thermo Fisher Scientific, MA, USA), then loaded into a shuttle under cryogenic conditions and transferred into an Aquilos dual-beam instrument equipped with a cryo-stage (FIB/SEM; Thermo Fisher Scientific). The sample surface was sputter-coated with platinum for 20 s at 30 mA current and then coated with a layer of organometallic platinum using the gas injection system for 6s at a distance of 1 mm before milling. Bulk milling was performed with a 30 kV gallium ion beam of 0.1 nA perpendicular to the grid on two side of a target yeast cell. The stage was then tilted to 10°-18° (so that the bulk-mill-holes lined up in front and behind the cell), and the cell was milled with 30 kV gallium ion beams of 100 pA current for rough milling and 30 pA for polishing until the final lamella was 100-200nm thick.

### Cryo-ET and image processing

Lamellae were imaged using a Titan Krios transmission electron microscope (FEI/Thermo Fisher Scientific) operated at 300 kV. Images were captured using a 4k × 4k K2 direct detection camera (Gatan, Pleasanton, CA) at a magnification of 26,000x (5.5 Å pixel size) or a 5k × 6k K3 direct detection camera (Gatan, Pleasanton, CA) at a magnification of 15,000x (5.7 Å pixel size). Tilt series were collected from 60° to -60° in 2° increments using a dose-symmetric tilting scheme (*32*). Counting modes of the K2 and K3 cameras were used and for each tilt image 15 frames (0.4 s exposure time per frame for K2 and 0.04 s exposure time per frame for K3) were recorded. Both cameras were placed behind a post-column energy filter (Gatan) that was operated in zero-loss mode (20-eV slit width). The defocus was set to -0.5 µm using a Volta phase plate (*33*). Data acquisition was performed using the microscope control software SerialEM (*34*) in low-dose mode, and the total electron dose per tilt series was limited to ∼100 e/Å^2^. The frames of each tilt series image were motion-corrected using MotionCor2 (*35*) and then merged using the script extracted from the IMOD software package (*36*) to generate the final tilt serial data set. Tilt series images were aligned fiducial-less using patch tracking (800 × 800-pixel size) and the tomogram was reconstructed by the back-projection method using the IMOD software package (*36*). To reduce noise, the cryo-tomograms were filtered with nonlinear anisotropic diffusion in the IMOD package.

### Radiolabeling, metabolite extraction, and metabolite separation

Approximately 100OD units of cells was used for radiolabeling experiments. All cells were grown and treated with AGR as previously described. Prior to radiolabeling, cells were collected by centrifugation and washed with dextrose-free media. All liquid was removed from cell pellets prior to labeling. To start radiolabeling, 1.5mL of dextrose-free media containing 5µCi/mL ^14^C-Acetate was quickly added to each tube, followed by mixing with pipetting. Tubes were tumbled in a 30C rotating incubator for fifteen minutes. To quench the radiolabeling reaction, and wash the cells, samples were pipetted into 40mL of -40C quenching buffer (60% methanol; 1mM tricine pH 7.4). Cells were incubated in quenching buffer for three minutes, centrifuged at 3,000xg at -10C, and washed with 20mL of -40C quenching buffer. Cells were again pelleted by centrifugation, and pellets for lipid extraction were stored at -80C. For pellets undergoing soluble metabolite extraction, all quenching buffer was thoroughly removed and 1.5mL of 80C 75% ethanol was added to each sample followed by a three minute incubation at 80C and a subsequent five minute incubation on ice. Debris was removed from metabolite extract by centrifuging at 20,000xg for one minute. Lipid extraction was performed as described above. HMG-CoA labeled during the pulse was separated and quantified as mevalonate. To convert endogenous HMG-CoA to mevalonate, ethanol was thoroughly evaporated from isolated metabolites under argon gas, and each sample was resuspended in HMGCR buffer (50mM Tris-HCl, pH 6.8; 100mM NaCl; 1mM MgCl_2_; 1mM DTT; 100mM glucose-6-phosphate; 1mM NADP+; 1mM NADPH). The samples were split into two tubes, one tube would be treated with enzymes, and the untreated tubes acted as blanks for endogenous mevalonate labeled during the pulse. For treated tubes, 2U of HMGCR enzyme (Sigma H7039) and 2U of glucose-6 phosphate dehydrogenase (Sigma G6378) were added. Reactions were carried out over night at 37C. Prior to loading samples onto TLC plates, total radioactivity in each sample was quantified by scintillation counting and loading was adjusted accordingly. HMGCR Reactions were spotted onto Silica gel G plates (Miles scientific P01911) and separated with a mobile phase of 70:25:5 Diethyl ether:glacial acetic acid:water. Each lane was doped with 5µg of cold mevalonate to act as a tracer for downstream scraping/quantification. TLC plates were developed overnight by autoradiography and visualized in an Amersham Typhoon FLA 9500 developer. To visualize the tracer mevalonate, plates were sprayed with p-anisaldehyde reagent and baked at 140C for ten minutes. Individual bands were scraped, mixed with 6mL of EcoLume scintillant, and quantified by scintillation counting in a Beckman LS 6500 instrument. Mevalonate from the untreated samples was averaged and subtracted from the final values of the treated samples. For lipid extracts, an aliquot of cell lysate was taken immediately following bead beating and radioactivity was quantified by scintillation counting to serve as a normalization standard prior to loading samples onto the TLC plate. To separate squalene, ergosterol, DAG, and sterol-esters, total lipid extracts were spotted onto silica gel 60G plates, and developed in a mobile phase of 55:35:10:1 hexane:petroleum ether:diethyl ether:glacial acetic acid. Autoradiography and scintillation counting was performed as indicated above. Prior to TLC separation, cold squalene, ergosterol and sterol-ester was added to each lane of the plate as tracer. DAG and ergosterol were quantified by densitometry in Fiji.

### Statistical analysis

T-tests and one-way ANOVA tests were performed using Graphpad Prism8 software. Kolmogorov-Smirnov test was conducted using kstest2 MATLAB function. All unpaired t-tests were performed with Welch’s correction. For one-way ANOVA, Brown-Forsyth and Welch ANOVA was performed to account for non-uniform standard deviations, followed by Turkey’s post-hoc test to extract p-values. For all t-tests and ANOVAs. * p-value<0.05, **p-value<0.01, ***p-value<0.001. Dotted lines in violin plots represent the median of the data, while upper and lower dotted lines represent the upper and lower quartiles. Bar graphs show mean of the data with error bars indicating the standard deviation.

## Acknowledgements

The authors want to thank members of the Henne and Nicastro labs for help in the completion of this project. The authors would also like to thank Joel Goodman for the *tgl3,4,5*,Δ yeast strain. We acknowledge Gang Fu and Evan Reetz for their technical assistance into the cryoFIB-SEM. W.M.H. is supported by funds from the Welch Foundation (I-1873), the NIH NIGMS (GM119768), and the UT Southwestern Endowed Scholars Program. S.R. is supported in part by a NIH T32 training grant (5T32GM008297). D.N. and L.G. are supported by grants from CPRIT (RP140082 to D.N.) The UTSW Cryo-EM Facility is supported in part by a CPRIT Core Facility Support Award RP170644.

---

**Table S1.**

| Strain | Description | Parent | Mat |
| --- | --- | --- | --- |
| YSR117 | Hmg1-GFP::NAT | W303 | a |
| YSR128 | Hmg1-GFP::NAT *nvj1*Δ | W303 | a |
| YSR136 | Hmg1-GFP::NAT *vac8*Δ | W303 | a |
| YSR12 | Hmg2-GFP::NAT | W303 | a |
| YSR134 | Hmg1<sub>1-525</sub>-GFP::NAT | W303 | a |
| YSR266 | *nvj1*Δ::HYG Hmg1-mRuby3::NAT<br>pRS305::ADH::Nvj1-mNeonGreen::LEU | W303 | a |
| YSR281 | *nvj1*Δ::HYG Hmg1-mRuby3::NAT<br>pRS305::ADH::Nvj1-PX::LEU | W303 | a |
| YSR282 | *nvj1*Δ::NAT Hmg1-mRuby3::HYG<br>pRS305::ADH::Nvj1<sub>Nvj2<sup>TM</sup></sub>-<br>mNeonGreen::LEU | W303 | a |
| YSR297 | pRS305::ADH::Nvj1<sub>15-24</sub>-<br>mNeonGreen | W303 | a |
| YSR298 | *nvj1*Δ::NAT Hmg1-mRuby3::HYG<br>pRS305::ADH::Nvj1<sub>15-30</sub>-<br>mNeonGreen | W303 | a |
| YSR214 | Hmg1-GFP::NAT<br>*upc2*Δ::HYG | W303 | a |
| YSR148 | *nvj1*Δ::HYG | W303 | a |
| ? | *tgl3,4,5*Δ | W303 | a |
| YSR176 | Hmg1-GFP::NAT<br>*osh1*Δ | W303 | a |
| YSR193 | Hmg1-GFP::NAT *ltc1*Δ | W303 | a |
| ? | Hmg1-GFP::NAT Nvj1-mRuby3::HYG | W303 | a |
| YSR149 | Hmg1-GFP::NAT<br>*snf1*Δ::HYG | W303 | a |
| YSR213 | Hmg1-GFP::NAT<br>*ecm22*Δ::HYG | W303 | a |
| YSR167 | Hmg1-mRuby3::HYG | W303 | a |
| YSR175 | Hmg1-mRuby3::HYG<br>*nvj1*Δ::NAT | W303 | a |
| YSR296 | Hmg1-DsRed::HYG | W303 | a |
| YSR301 | Hmg1-DsRed::HYG<br>*nvj1*Δ::NAT | W303 | a |
| YSR250 | *hmg1*Δ::HYG<br>*hmg2*Δ::NAT<br>pRS305::ADH::Hmg1-GFP::LEU | W303 | a |
| YSR260 | *hmg1* Δ::HYG<br>*hmg2* Δ::NAT *nvj1Δ::HIS*<br>*pRS305::ADH::Hmg1-*<br>*GFP::LEU* | W303 | a |
| YSR183 | Erg10-mNeonGreen::HYG | W303 | a |
| YSR160 | Erg13-mNeonGreen::HYG | W303 | a |
| YSR159 | Erg12-mNeonGreen::HYG | W303 | a |
| YSR187 | Erg8-mNeonGreen::HYG | W303 | a |
| YSR162 | Mvd1-mNeonGreen::HYG | W303 | a |
| YSR161 | Idi1-mNeonGreen::HYG | W303 | a |
| YSR188 | Erg9-mNeonGreen::HYG | W303 | a |
| YSR156 | Erg1-mNeonGreen::HYG | W303 | a |
| YSR186 | Erg7-mNeonGreen::HYG | W303 | a |
| YSR184 | Erg11-mNeonGreen::HYG | W303 | a |
| YSR276 | Ncp1-mNeonGreen::HYG | W303 | a |
| YSR185 | Erg24-mNeonGreen::HYG | W303 | a |
| YSR182 | Erg25-mNeonGreen::HYG | W303 | a |
| YSR251 | Erg26-mNeonGreen::HYG | W303 | a |
| YSR236 | Erg27-mNeonGreen::HYG | W303 | a |
| YSR275 | Erg28-mNeonGreen::HYG | W303 | a |
| YSR270 | Erg6-mNeonGreen::HYG | W303 | a |
| YSR157 | Erg2-mNeonGreen::HYG | W303 | a |
| YSR189 | Erg5-mNeonGreen::HYG | W303 | a |
| YSR158 | Erg4-mNeonGreen::HYG | W303 | a |
| PK1359 | Erg6-mTFP::HIS3<br>Whi5-mKok::TRP1<br>Vma1-mNeptune2.5::KAN<br>Nvj1-mRuby3::NAT<br>Msn2-mNeonGreen::HYG | W303 | Diploid |
| PK1261 | Erg6-mTFP::HIS3<br>Whi5-mKok::TRP1<br>Vma1-mNeptune2.5::KAN<br>nvj1Δ::NAT<br>Msn2-mNeonGreen::HYG | W303 | Diploid |

**Table S2.**
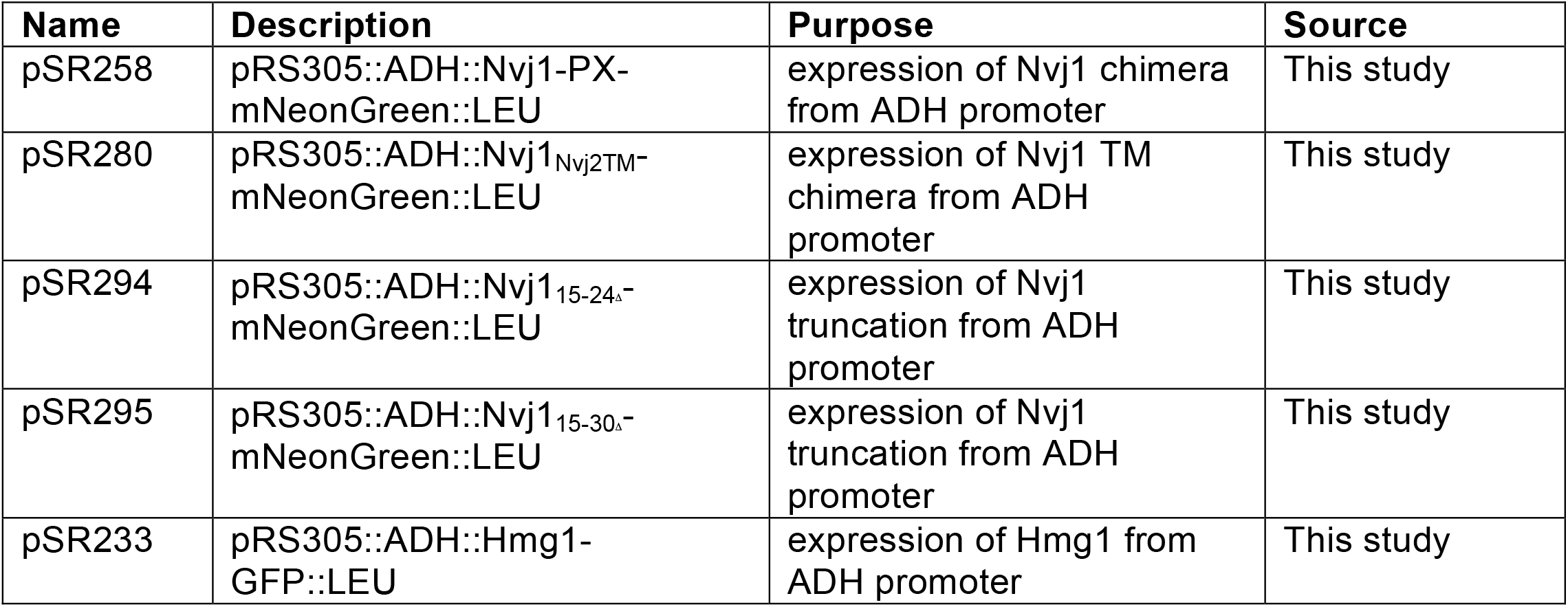

**Fig S1.**
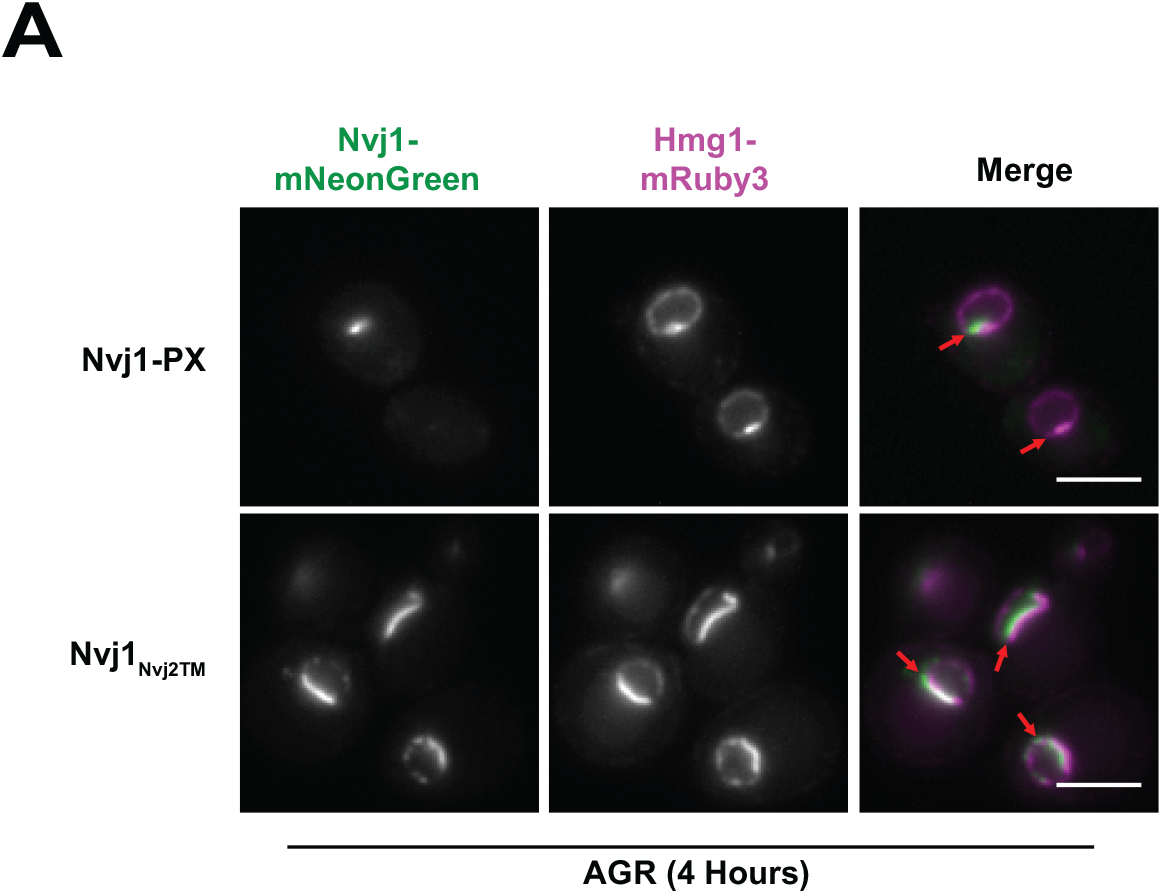
Hmg1 NVJ partitioning is not dependent on the cytoplasmic or transmembrane regions of Nvj1. (A) Epifluorescence microscopy images of cells expressing ADH promoter-driven Nvj1-mNeonGreen (Nvj1-mNg) chimeric constructs (green) either substituting the endogenous transmembrane region with that of Nvj2 (Nvj1NVJ2TM) or substituting the endogenous Vac8 binding domain with a vacuole binding PX domain (Nvj1PX). Chimera strains co-expressed endogenously tagged Hmg1-mRuby3 (magenta) and lacked endogenous Nvj1. Images were taken after exposing cells to four hours of AGR. Scale bars represent 5mm.

**Fig S2.**
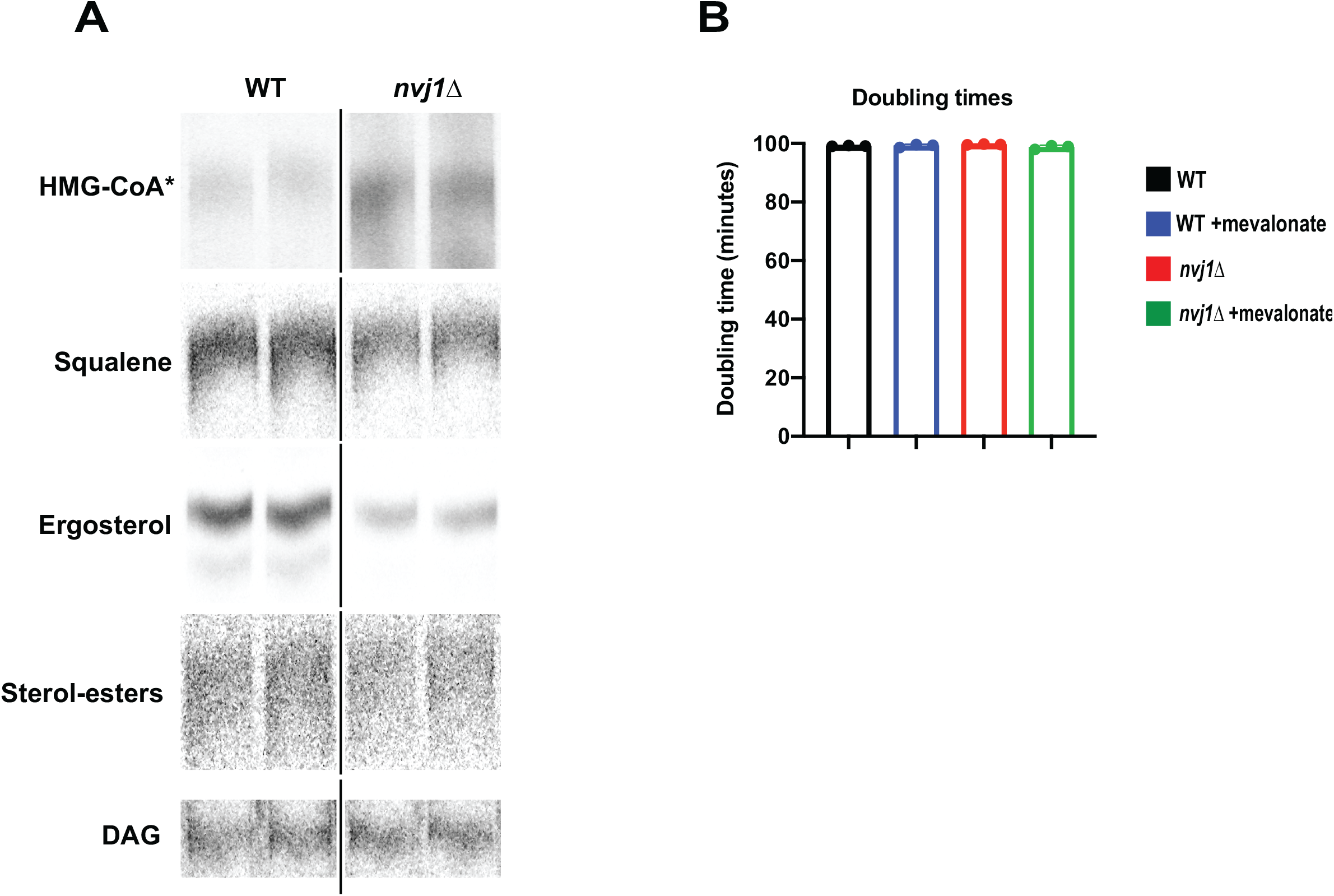
Cells lacking endogenous Nvj1 accumulate Hmg1 substrates and have decreased mevalonate-derived pathway products. (A) Autoradiograms taken of TLC plates to visualize HMG-CoA* (visualized as mevalonate, see Fig 3 legend or supplemental methods) or squalene, ergosterol, SE, and DAG in wild type (WT) or *nvj1*Δ cells. Squalene, ergosterol, SE, and DAG were visualized on the same plate. DAG was used as a control band for neutral lipid species. Brightness/contrast was adjusted linearly in ImageJ to visualize bands. (B) Doubling times of cells from Fig 5H. (Brown-Forsyth and Welch ANOVA. N=3).

**Fig S3.**
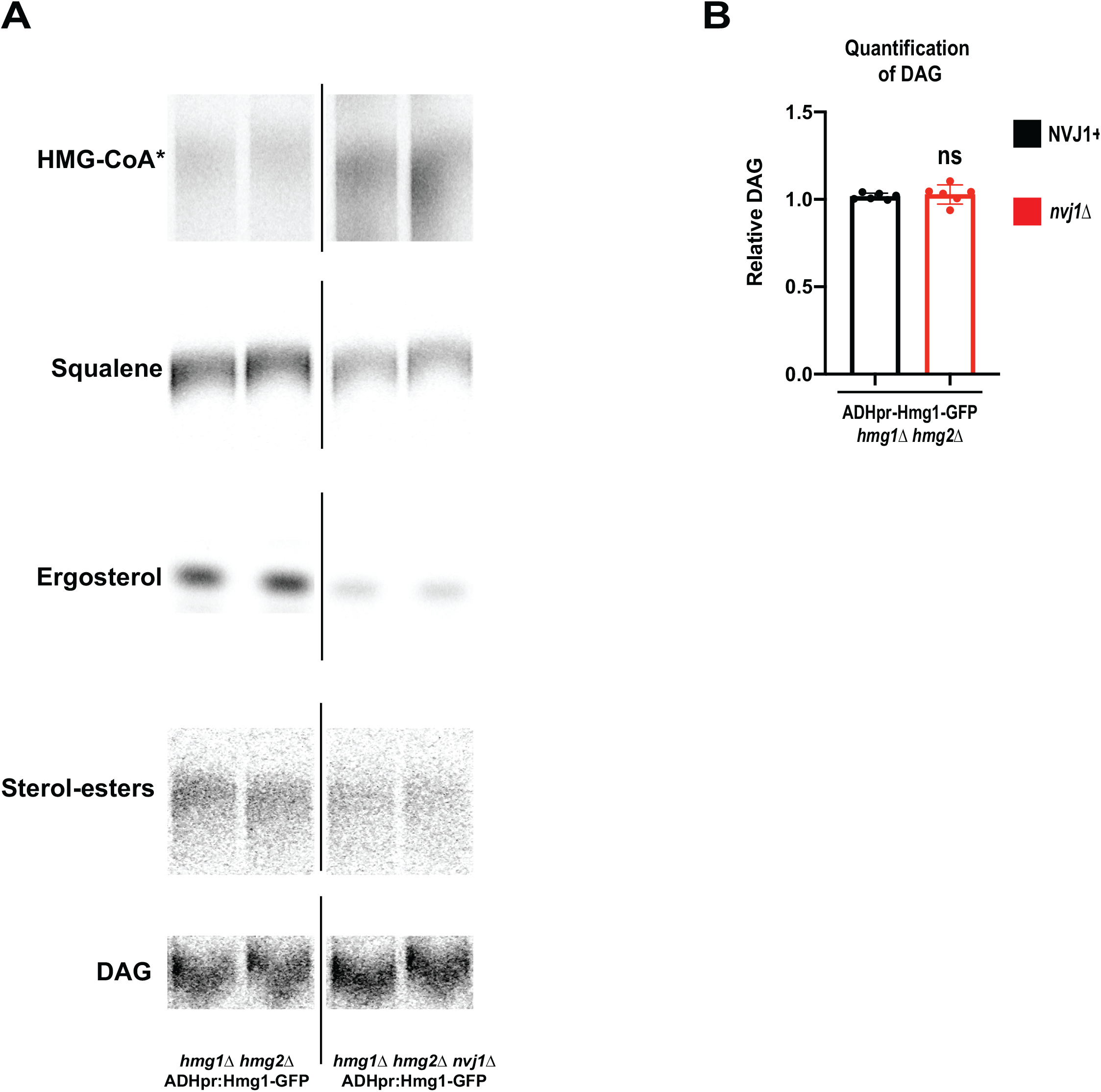
Cells expressing ADHpr:Hmg1-GFP in the absence of endogenous Nvj1 have exacerbated Hmg1 catalytic deficiency phenotype. (A) Autoradiograms taken of TLC plates to visualize HMG-CoA* (as previously) or squalene, ergosterol, SE, and DAG in ADHpr:Hmg1-GFP lines in the presence or absence of Nvj1. Bands were visualized as described in Fig S5. (B) Densitometry quantification of DAG bands as shown in (A). (Brown-Forsyth and Welch ANOVA. N=6).

**Fig S4.**
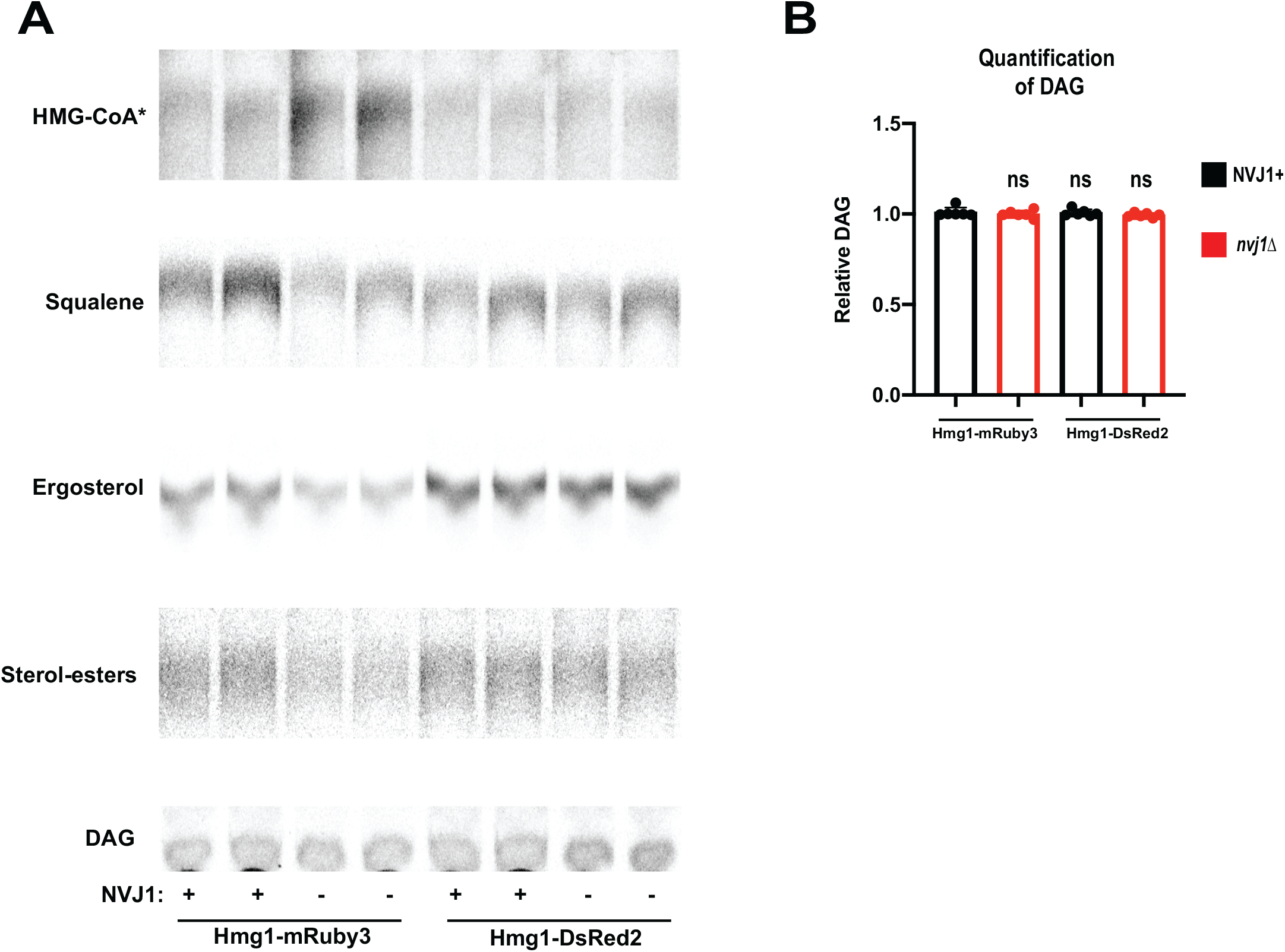
Artificial inter-enzyme association via DsRed2 tagging rescues *nvj1*Δ associated Hmg1 catalytic deficiency. (A) Autoradiograms taken of TLC plates to visualize HMG-CoA* (as previously described) or squalene, ergosterol, SE, and DAG in cells expressing endogenously tagged monomeric Hmg1-mRuby3 or tetrameric Hmg1-DsRed2 in the presence or absence of endogenous Nvj1. Bands were visualized as described in Fig S3. (B) Densitometry quantification of DAG bands as shown in (A). (Brown-Forsyth and Welch ANOVA. N=6).

**Fig S5.** AGR promotes the core of LDs to bubble under high EM electron doses. (A-E) 2D cryo-EM images of the same LD from a cryo-FIB milled WT yeast cell cultured with glucose after applying an increasing amount of electron dose (1 - 400 e-/Å2); note that no liquid-crystalline layers were observed and bubbling did not occur even at a high electron dose when cytoplasmic structures exhibited bubbling (arrowheads) at 400 e-/Å2. (F-J) 2D cryo-EM images of the same 2 LDs from a cryo-FIB milled WT yeast cell cultured in AGR after applying an increasing amount of electron dose (1 - 161 e-/Å2); note that liquid-crystalline layers are clearly visible in the LDs (the boxed area in G is magnified in J), and that radiation damage - seen as white bubbling - becomes clearly visible in the layer-free center of the LDs starting at a dose between 20-30 e-/Å2. The black density in the top-left corner is ice contamination that shifted position during beam exposure (beam-induced motion). (J) Enlarged area from (G), showing the peripheral, concentric LC-layers of the LD. Scale bar, 50 nm (A-I) and 25 nm (J).

