## Supplementary figures and images for "Glucose restriction drives spatial re-organization of mevalonate metabolism and liquid-crystalline lipid droplet biogenesis"

### Supp Figure 5

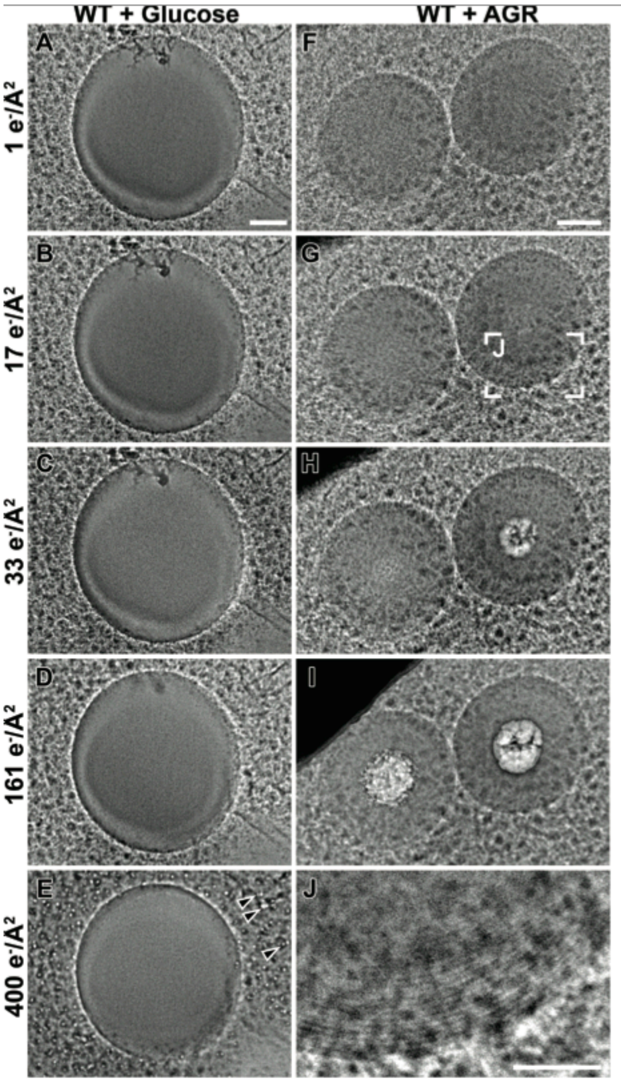
